# An increased excitation and inhibition onto CA1 pyramidal cells sets the path to Alzheimer’s disease

**DOI:** 10.1101/2025.05.27.656417

**Authors:** Patrick H. Wehrle, Travis J. Rathwell, Maurice A. Petroccione, Ethan D. Caiazza, Anthony K. Manning, Leonardo Frasson dos Reis, Gabrielle C. Todd, Nurat Affinnih, Saad Ahmad, Hasan Mehdi, Ian L. Tschang, Umair Hassan, Brianna R. Tsakh, Marisol C. Lauffer, Paul A. Rosenberg, Martin Darvas, David G. Cook, Annalisa Scimemi

## Abstract

Synapses are critical targets of Alzheimer’s disease (AD), a highly prevalent neurodegenerative disease associated with accumulation of extracellular amyloid-β peptides. Although amyloidosis and aggregation of the 42-amino acid amyloid-β (Aβ_42_) have long been considered pathogenic triggers for AD, clinical evidence linking high levels of Aβ_42_ with normal cognition challenges this hypothesis. To resolve this conundrum on the role of Aβ_42_ in regulating synaptic activity, we used an adeno-associated viral vector approach that triggers extracellular accumulation of Aβ_42_ and spatial memory impairment. We show that Aβ_42_ leads to an early increase in excitatory and proximal inhibitory synaptic transmission onto hippocampal CA1 pyramidal cells, and an increased expression of the glutamate transporter GLT-1 in these cells. Aβ_42_ accumulation does not cause early cognitive deficits unless accompanied by an increased neuronal GLT-1 expression, suggesting this transporter is a critical mediator of Aβ_42_’s effects. These findings unveil key molecular and cellular mechanisms implicated with AD pathogenesis.

## INTRODUCTION

AD is one of the most prevalent and impactful neurodegenerative diseases, the fifth-leading cause of death among Americans aged 65 and older and accounts for up to two third of dementia cases^1,2^. The strongest predictor of cognitive decline in AD is synaptic loss and neuronal death^3^. Although extensive neuronal loss and astrogliosis can be detected post-mortem in the neocortex of AD patients, multiple lines of evidence indicate that neuronal hyperactivity precedes neuronal loss in the early stages of AD. For example, functional MRI studies in patients with mild cognitive impairment (MCI) and in pre-symptomatic carriers of familial AD detect increased activity in the neocortex^4–13^. Consistent with these findings, studies using mouse models of AD show that neuronal hyperactivity occurs early and is pronounced in AD-vulnerable brain regions implicated with memory and cognition, like the hippocampus^5,14–18^. In addition, there is an increased risk for epilepsy and seizures in AD patients that is greater at earlier stages of the disease^19,20^. This increased neuronal activity has been suggested to be a critical driving factor for the development and progression of AD, as it could increase extracellular levels of Aβ peptides and speed AD pathology^21–27^.

Deposition of Aβ in brain regions that control memory and cognition is considered essential for the neuropathologic diagnosis of AD^28,29^. Once applied to acute rat hippocampal slices, soluble Aβ oligomers isolated from AD patients decrease synaptic function and, *in vivo,* they impair memory recall^28,30–32^. Among all known Aβ peptides, Aβ_42_ represents the earliest form to accumulate in the brain and the most abundant protein in neuronal plaques of AD patients, suggesting this peptide could be critical for AD pathology^33^. However, clinical trials aiming to reduce Aβ_42_ production and aggregation and to promote Aβ_42_ clearance had mixed or only moderate results for cognitive or functional benefits^34,35^. Obviously one major dilemma is whether any of these strategies could be more successful if started before any damage occurs, but without tools for an early diagnosis of AD, this remains an unsurmountable challenge. Because of these apparently conflicting findings, the question of whether Aβ_42_ has a function in regulating neocortical activity and AD progression remains more contentious and debated than ever.

Currently available transgenic models of AD rely on overexpression of mutant human amyloid precursor protein and PS1, but the expression of many different cytotoxic proteolytic peptides is increased in these models, making it difficult to understand the specific role of Aβ_42_^36–39^. To overcome these limitations, we used an adeno-associated viral (AAV) vector approach to trigger extracellular accumulation of Aβ_42_ in the mouse hippocampus. This strategy is similar to that used by Lawlor et al. which showed that AAV-BRI-Aβ_42_ is sufficient to initiate deposition of insoluble Aβ in diffuse plaque-like structures 3 months post-injection into the hippocampus^40^. Here, we analyze the effect of AAV-Aβ_42_ earlier on, 3-8 weeks post injections, to identify the early effects of this peptide, including those of Aβ_42_ that is not yet aggregated in plaques on excitatory and inhibitory transmission onto CA1 pyramidal cells (CA1-PCs). Our analysis is not limited to neurons, but also astrocytes, as these cells play a critical role in regulating brain homeostasis and synaptic activity in the healthy brain, and have also been suggested to contribute to the neuropathologic process of AD^41^.

## RESULTS

### AAV-Aβ_42_ induces Aβ_42_ accumulation in hippocampal area CA1

We used two AAV vectors in this work, which we refer to as AAV-Sham and AAV-Aβ_42_. For simplicity, we refer to mice that received stereotaxic injections of one or the other AAV as Sham and Aβ_42_ mice. The AAVs were injected into the CA1-PC layer (**Fig.1A**), and we verified that they transfected CA1-PCs using confocal imaging experiments on tissue samples collected 3-8 weeks post injection (**Fig.1B**). This time window was sufficient to disrupt spatial learning, as indicated by subjecting mice to a novel object location task (**Fig. 1C**). In this test, we first habituated mice to an empty open field arena, then we trained them to recognize two identical objects located in adjacent corners of the field. Last, after 90 min, we displaced one of the two objects to the opposite corner of the field and asked whether the mice spent more time around the displaced or the non-displaced object. C57BL/6J wild type (WT) and Sham mice had a place preference for the displaced object during the testing phase (**Fig. 1C**). By contrast, the Aβ_42_ mice spent a similar proportion of time near the two objects (**Fig. 1C**). We confirmed using Mesoscale Discovery (MSD) assays that 3-8 weeks post AAV-Aβ_42_ injections there was an increased level of detergent soluble (**Fig. 1D, left**) and insoluble Aβ_42_ in the hippocampus (**Fig. 1D, right**). Consistent with these findings, an increase in Aβ_42_ immunoreactivity, with no plaques, was detected in hippocampal area CA1 of Aβ_42_ mice, not in naïve WT or Sham mice (**Fig. 1E**). Although Aβ_42_ is a trigger of tau protein phosphorylation, we did not detect a significant immunoreactivity for the phospho-tau Ser262 (**Fig. 1F**)^42^. In AD, there is a significant activation of proteins within the caspase family, known to mediate apoptotic signaling pathways and neurodegeneration^43–46^. For this reason, we asked whether AAV-induced Aβ_42_ expression enhanced the expression of activated caspase-3, resulting from caspase cleavage at Asp175 (**Supp.** Fig. 1). However, we did not detect caspase-3 immunolabeling in sections from WT, Sham and Aβ_42_. The only detectable staining was obtained in mouse hippocampal sections maintained for 30 min without oxygen supply, which we used as a positive control for our antibody staining (**Supp.** Fig. 1). Together, these findings indicate that AAV-Aβ_42_ is effective at boosting Aβ_42_ protein accumulation without inducing plaque formation and cell death in the mouse hippocampus, and at triggering hippocampal-dependent cognitive deficits.

**Figure 1.**
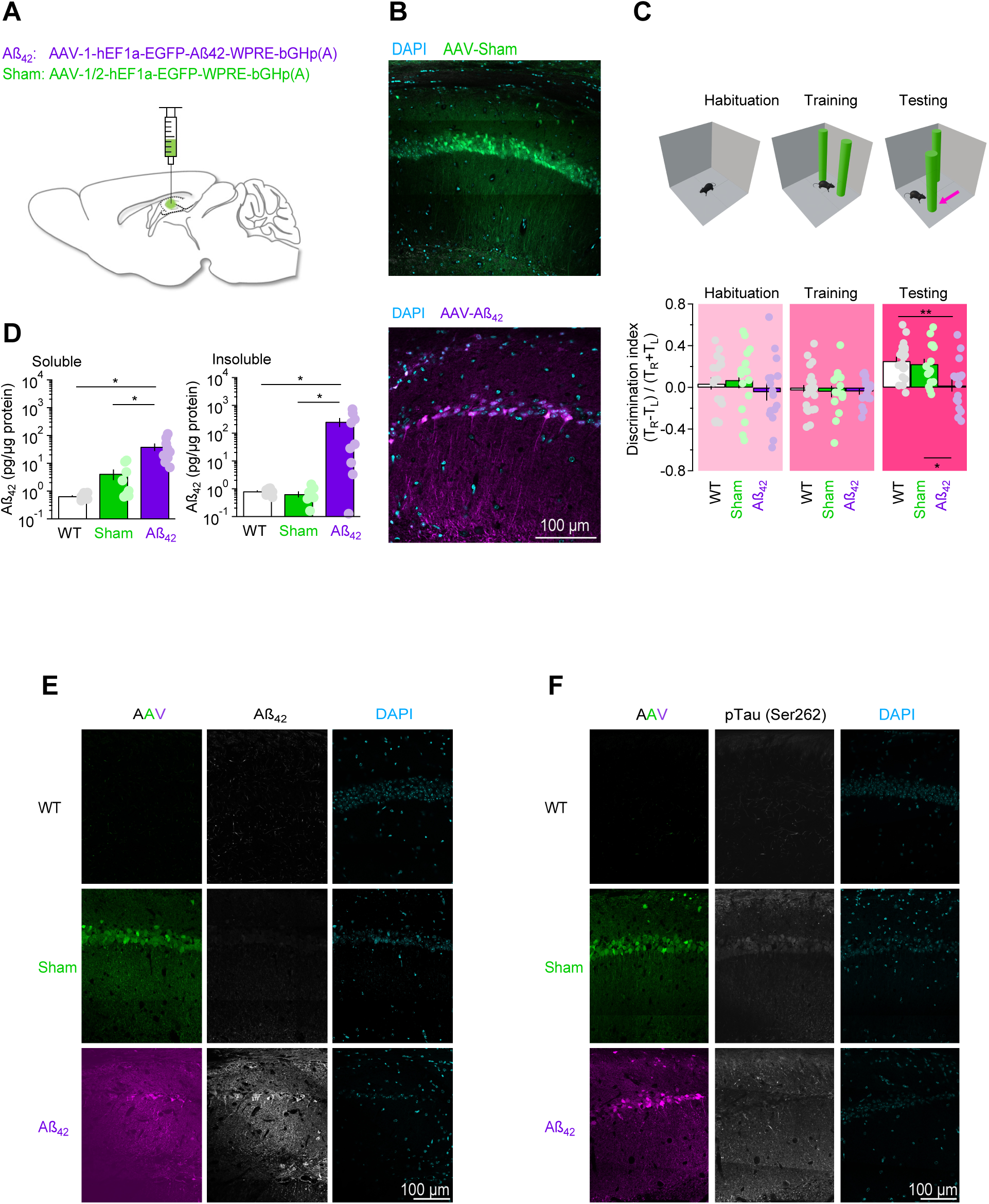
Functional characterization of AAV-Sham and AAV-Aβ_42_. **(A)** Schematic representation of the stereotaxic injections used to deliver AAV-Sham or AAV-Aβ_42_ to hippocampal area CA1 of P14-16 mice. **(B)** Confocal images of the mouse hippocampus 3 weeks after stereotaxic injection of 200 nl AAV-Sham (*top*) or AAV-Aβ_42_ (*bottom*) in hippocampal area CA1. **(C)** Summary of object location test, showing that Aβ_42_ mice do not discriminate the location of a displaced object (WT, *N*=12, n=20; Sham, *N*=14, n=16, Aβ_42_, *N*=16, n=16). One-way ANOVA followed by pairwise comparisons. **(D)** Radio-immunoprecipitation assay (RIPA) for detection of soluble (*left*) and insoluble (*right*) Aβ_42_ in WT (*white, N=12*), Sham (*green, N=8*) and Aβ_42_ mice (*purple, N=9*). Mann-Whitney test. **(E)** Immunofluorescence labeling of hippocampal sections from WT (*top*), Sham (*middle*) and Aβ_42_ mice (*bottom*) using anti-EFGP (*left*) and anti-Aβ_42_ antibodies (*center*). DAPI nuclear staining is shown in cyan (*right*). **(F)** As in E, using anti-EFGP (*left*) and anti-phosphoprotein tau antibodies (*center*). Data represent mean±SEM. \**p*<0.05; ** *p*<0.01; \*\*\**p*<0.001.

### Aβ_42_ does not change the gross morphology but increases spine density of CA1-PCs

To determine whether Aβ_42_ altered the overall structure of CA1-PCs, we obtained biocytin fills from CA1-PCs of WT, Sham and Aβ_42_ mice (**Supp.** Fig. 2A,B). A Sholl analysis showed that the morphology of these cells was indistinguishable across the three groups of mice (**Supp.** Fig. 2C). The cumulative distribution of the branch intersections with circles of increasing radii, centered at the soma, was similar for CA1-PCs of WT, Sham and Aβ_42_ mice (**Supp.** Fig. 2D). Consistent with these findings, we did not detect significant differences in the total and average number of intersections, and in the maximum radius of the biocytin filled CA1-PCs (**Supp.** Fig. 2E-G).

Upon closer inspection, we noticed that the apical dendrites of CA1-PCs were studded with more spines in Aβ_42_ compared to WT and Sham mice (**Fig. 2A,B**). Given that the percentage distribution of mushroom, thin and stubby spines was not different in these three cohorts of mice, we believe that Aβ_42_ induces a proportionate increase in the linear density of all these different types of spines (**Fig. 2C**). On average, the surface area, volume, head diameter and neck length of spines in each class was not different between WT, Sham and Aβ_42_ mice (**Fig. 2D-G**). We hypothesized that an increased spine density could lead to an increased frequency of action potential independent miniature EPSCs (mEPSCs), which we detected experimentally in CA1-PCs voltage clamped at −70 mV, in the presence of picrotoxin (100 µM), a GABA_A_ receptor antagonist, in WT, Sham and Aβ_42_ mice (**Fig. 2H,I**). At this holding potential, the mEPSC are mostly mediated by AMPA receptors. The mEPSC amplitude and kinetics were comparable across the three groups, suggesting that Aβ_42_ does not change the quantal size of glutamatergic events recorded from CA1-PCs (**Fig. 2H-K**). Although the increased in mEPSC frequency could be induced by changes in presynaptic release probability (P_r_), the experiments described in **Fig. 3** allowed us to rule out this hypothesis.

**Figure 2.**
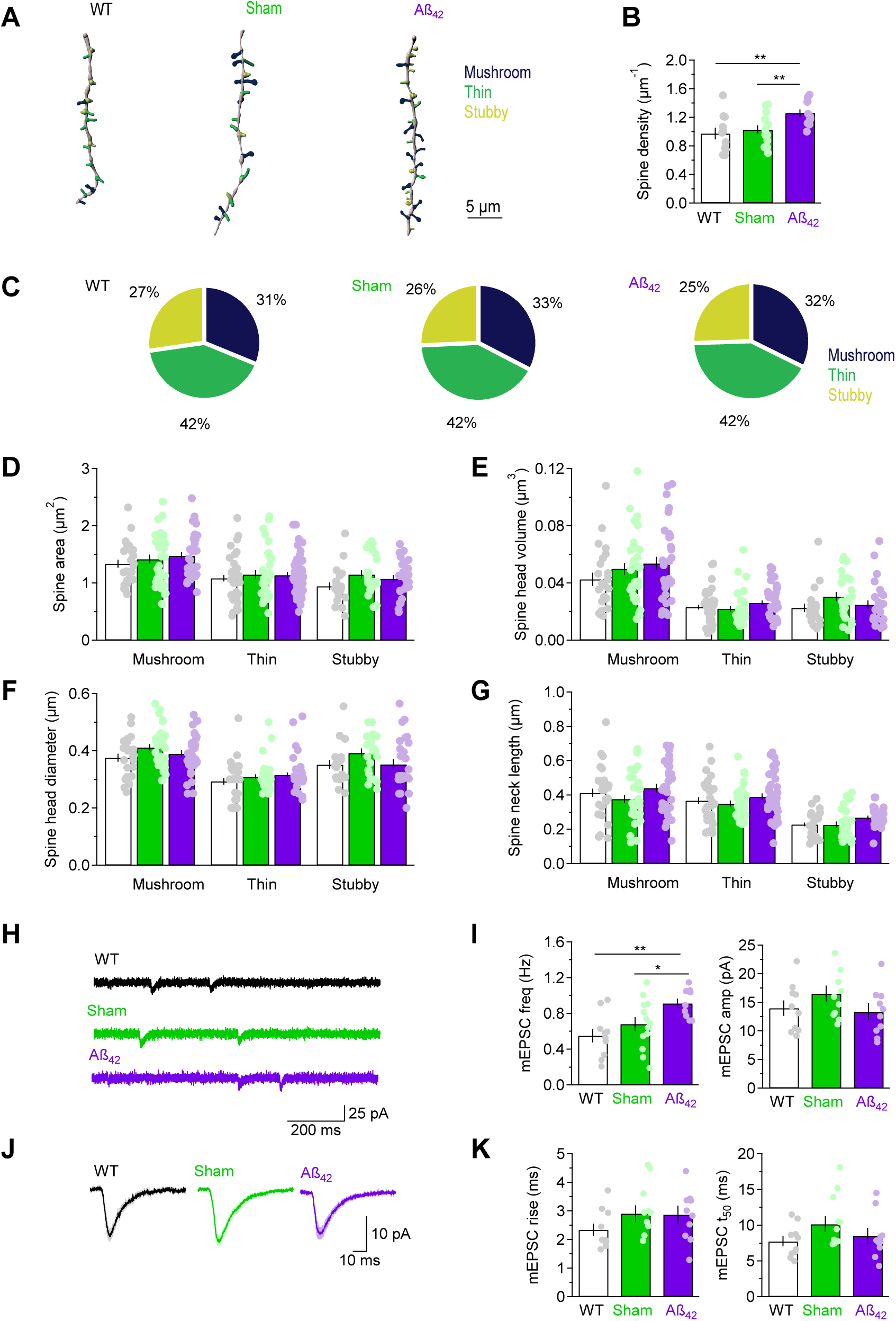
AAV-Aß_42_ leads to an early-onset increase in spine density and mEPSC frequency in CA1-PCs. **(A)** Example of 3D reconstructions of dendritic branches of CA1-PCs. Mushroom, thin and stubby spines are color coded in *black*, *green*, and *olive*. **(B)** Summary of the spine density in CA1-PCs of WT (*N*=4, n=11), Sham (*N*=5, n=13) and Aβ_42_ mice (*N*=4, n=11). **(C)** Pie charts showing the percentage distribution of mushroom, thin and stubby spines in CA1-PCs of WT (*N*=4, n=24, 32, 21), Sham (*N*=5, n=33, 34, 23) and Aβ_42_ mice (*N*=4, n=33, 43, 26). **(D)** Summary of the surface area of different types of spines in WT (*white*), Sham (*green*) and Aβ_42_ mice (*purple*). **(E-G)** As in D, for spine head volume (E), diameter (F) and neck length (G). **(H)** Example of mEPSC recordings in CA1-PCs of WT, Sham, and Aβ_42_ mice voltage clamped at −70 mV. **(I)** Summary of mEPSC frequency (*left*) and amplitude (*right*) for WT (*N*=7, n=10), Sham (*N*=8, n=13), and Aβ_42_ (*N*=4, n=9). **(J)** Average mEPSCs recorded in CA1-PCs of WT, Sham, and Aβ_42_ mice. **(K)** As in I, for 20-80% rise time (*left*) and 50% decay time (*right*). Data represent mean±SEM. One-way ANOVA followed by pairwise comparisons. \**p*<0.05; ** *p*<0.01; \*\*\**p*<0.001.

**Figure 3.**
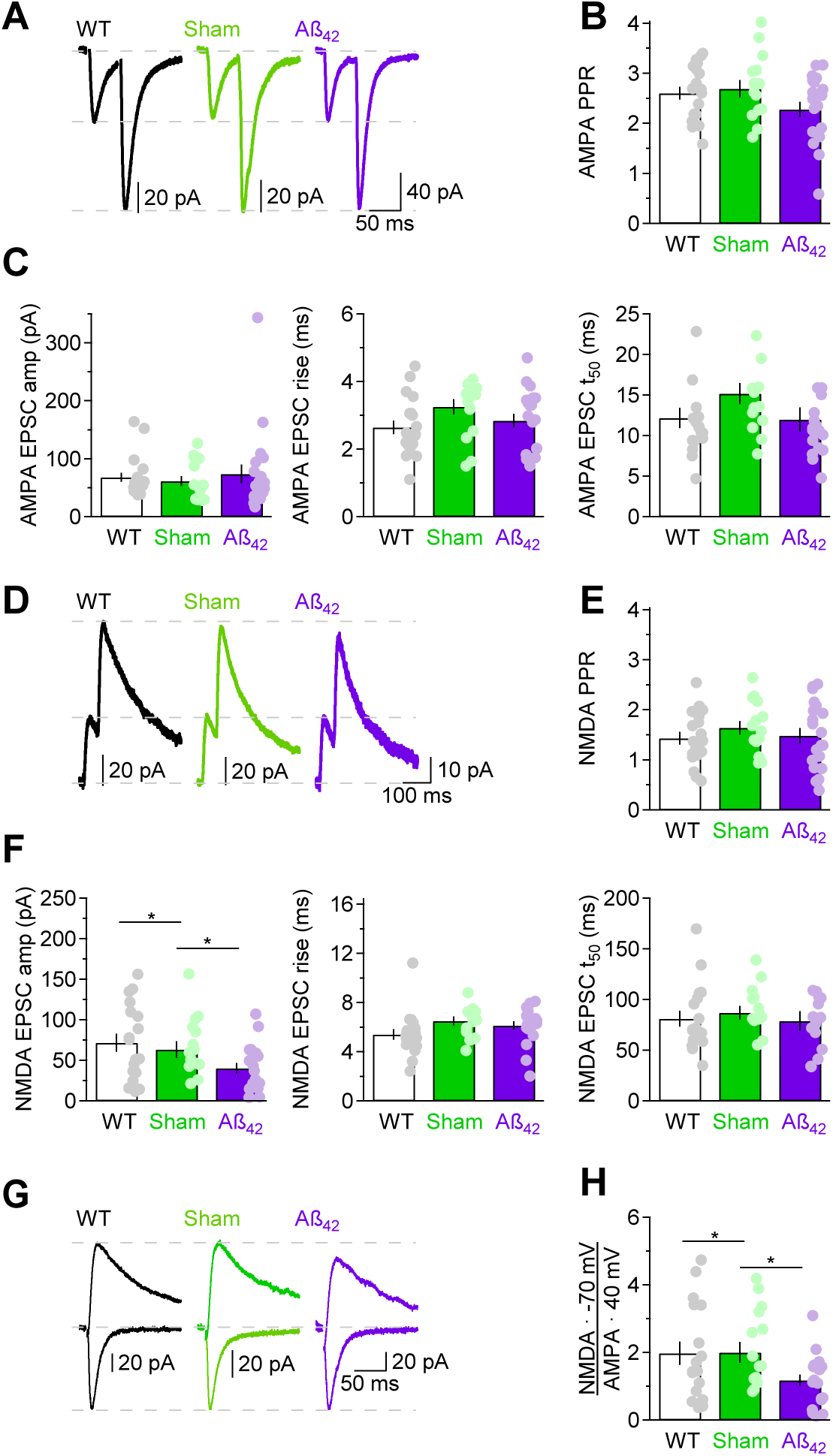
AAV-Aß_42_ leads to a reduction in the NMDA receptor activation. **(A)** Representative paired AMPA EPSCs evoked by electrical stimulation of Schaffer collaterals in WT, Sham, and Aβ_42_ mice. The recordings were obtained from CA1-PCs voltage clamped at −70 mV, in the presence of the GABA_A_ receptor antagonist picrotoxin (100 µM). Each trace represents the average of 20 EPSCs. **(B)** Summary of the paired pulse ratio of AMPA EPSCs in WT (*N*=15, n=19), Sham (*N*=12, n=15) and Aβ_42_ mice (*N*=14, n=20). **(C)** The amplitude (*left*), 20-80% rise (*center*) and 50% decay time (*right*) of electrically evoked AMPA EPSCs is similar in WT, Sham and Aβ_42_ mice. **(D-F)** As in A-C, for NMDA EPSCs recorded in a subset of CA1-PCs voltage clamped at 40 mV, in the presence of picrotoxin (100 µM) and NBQX (10 µM) in WT (*N*=15, n=18), Sham (*N*=12, n=15) and Aβ_42_ mice (*N*=14, n=20). **(G)** In-cell comparison of AMPA and NMDA EPSCs. The purple trace shows that the NMDA/AMPA ratio is reduced in CA1-PCs from Aβ_42_ mice. The traces are scaled with respect to the peak of the AMPA EPSCs. **(H)** Summary of the NMDA/AMPA EPSC amplitude ratio corrected by the driving force of glutamatergic currents, with an estimated reversal potential of 0 mV,) in WT (*N*=15, n=17), Sham (*N*=12, n=15) and Aβ_42_ mice (*N*=14, n=20). The NMDA/AMPA ratio is significantly reduced in CA1-PCs from Aβ_42_ mice. Data represent mean±SEM. One-way ANOVA followed by pairwise comparisons. \**p*<0.05; ** *p*<0.01; \*\*\**p*<0.001.

### Aβ_42_ reduces NMDA receptor activation, not the number of functional NMDA receptors in CA1-PCs

To determine whether Aβ_42_ induces changes in P_r_, we delivered single and paired stimuli to Schaffer collaterals in *stratum radiatum* (*s.r.*) and recorded electrically evoked EPSCs (eEPSCs) in CA1-PCs (**Fig. 3A**). The stimulus intensity was set to evoke eEPSCs of similar amplitude in CA1-PCs of WT, Sham and Aβ_42_ mice. The amplitude ratio between the second and first eEPSCs, known as the paired-pulse ratio (PPR), is inversely proportional to P_r_. In our experiments, the PPR, amplitude, rise and decay time of AMPA eEPSCs recorded at −70 mV was similar in WT, Sham and Aβ_42_ mice, suggesting that P_r_ and AMPA receptor activation is not altered by Aβ_42_ (**Fig. 3B,C**). We also calculated the PPR of NMDA eEPSCs recorded at a holding potential of 40 mV, in the presence of AMPA and GABA_A_ receptor antagonists NBQX (10 µM) and picrotoxin (100 µM), respectively (**Fig. 3D,E**). The PPR of NMDA eEPSCs was similar across the three mouse groups, as were the NMDA eEPSC rise and decay times (**Fig. 3F**). However, the amplitude of the NMDA eEPSCs was smaller in Aβ_42_ compared to WT and Sham mice (**Fig. 3F**). Consistent with these findings, an in-cell comparison of AMPA and NMDA eEPSCs showed that the NMDA/AMPA eEPSC ratio, corrected by the driving force for each receptor, was reduced in Aβ_42_ compared to WT and Sham mice (**Fig. 3G,H**). This finding suggests that the number and/or activation of NMDA receptors might be reduced in response to an increased extracellular concentration of Aβ_42_, a hypothesis that we tested with additional experiments.

To determine whether the reduced NMDA receptor activation in Aβ_42_ mice was due to a reduced plasma membrane expression of NMDA receptors, we performed a series of flash uncaging experiments in which RuBi-glutamate (50 µM) was added to the recording solution, and 5 ms flashes of blue light (∼250 µW at the sample plane) were delivered to the whole field of view of our microscope, to cover the entire dendritic arborization of the cell we patched (**Fig. 4A, left**). The intensity of the light pulses was the same for all recordings included in this dataset. We first recorded AMPA uncaging EPSCs (uEPSCs) in CA1-PCs voltage clamped at −70 mV, in the presence of picrotoxin (100 µM; **Fig. 4A, right**). The amplitude, rise and decay of AMPA uEPSCs was similar in WT, Sham and Aβ_42_ mice, suggesting that Aβ_42_ does not change the functional pool of AMPA receptors in CA1-PCs (**Fig. 4B**). In a subset of cells, we blocked AMPA receptors with NBQX (10 µM), switched the holding potential to 40 mV and recorded NMDA uEPSCs (**Fig. 4C**). The amplitude, rise and decay of NMDA EPSCs was also similar in WT Sham and Aβ_42_ mice, suggesting that Aβ_42_ does not change the functional pool of NMDA receptors in CA1-PCs (**Fig. 4D**). As a result, the NMDA/AMPA ratio of uEPSCs was similar in WT, Sham and Aβ_42_ mice (**Fig. 4E, F**). Based on these findings, we infer that the reduced activation of NMDA receptors detected when recording eEPSCs is not due to an Aβ_42_-induced reduction in the plasma membrane expression of NMDA receptors, but rather to a reduced recruitment of these receptors by synaptically-released glutamate in Aβ_42_ mice. Because the functional pool of AMPA and NMDA receptors is similar in WT, Sham, and Aβ_42_ mice, but there are more excitatory synapses in CA1-PCs of Aβ_42_ mice, we suggest this might be due to Aβ_42_ promoting the recruitment of extrasynaptic NMDA receptors to synaptic sites, via subcellular redistribution of these receptors on the plasma membrane of CA1-PCs.

**Figure 4.**
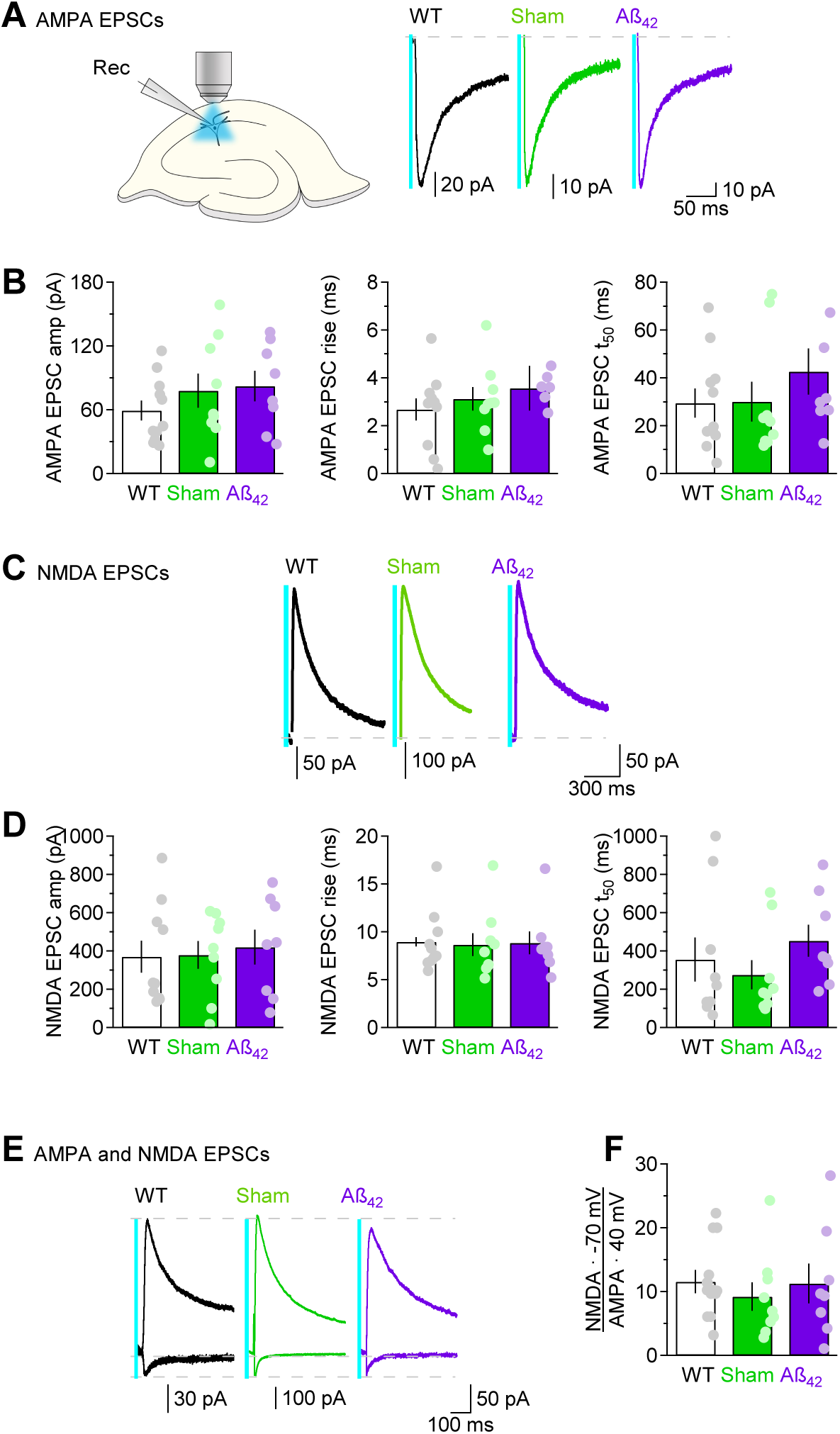
CA1-PCs have a similar pool of functional AMPA and NMDA receptors in WT, Sham and Aβ_42_ mice. **(A)** *Left,* Schematic representation of EPSCs evoked by delivering full-field, 5 ms-long blue light pulses to slices perfused with RuBi-glutamate (50 µM). The light intensity was maintained constant across slices from WT, Sham and Aβ_42_ mice. *Right,* Representative oEPSCs evoked in CA1-PCs of WT, Sham and Aβ_42_ mice. Each trace represents the average of 10 oIPSCs. The vertical cyan line represents the artifact of the optical stimulation. **(B)** Summary of the AMPA oEPSC amplitude (*left*) and kinetics (*center, right*) recorded in CA1-PCs voltage clamped at −70 mV, in the presence of picrotoxin (100 µM), in WT (*N*=9, n=11), Sham (*N*=8, n=9) and Aβ_42_ mice (*N*=7, n=8). **(C)** In a subset of CA1-PCs, we recorded NMDA oEPSCs, without changing the stimulus intensity. The traces represent NMDA oEPSCs recorded from CA1-PCs of WT, Sham and Aβ_42_ mice, voltage clamped at 40 mV and in the presence of picrotoxin (100 µM) and NBQX (10 µM). Each trace represents the average of 10 oIPSCs. The vertical cyan line represents the artifact of the optical stimulation. **(D)** Summary of the NMDA oEPSC amplitude (*left*) and kinetics (*center, right*), in WT (*N*=9, n=10), Sham (*N*=8, n=9) and Aβ_42_ mice (*N*=7, n=8). **(E)** In-cell comparison of AMPA and NMDA oEPSCs recorded from CA1-PCs of WT (*N*=9, n=10), Sham (*N*=8, n=9) and Aβ_42_ mice (*N*=7, n=8). **(F)** The NMDA/AMPA ratio is similar across the three different cohorts of mice, suggesting that the functional pool of AMPA and NMDA receptors is not altered by AAV-Aβ_42_. Data represent mean±SEM. One-way ANOVA followed by pairwise comparisons. \**p*<0.05; ** *p*<0.01; \*\*\**p*<0.001.

### Fewer astrocytes take up glutamate, yet glutamate clearance is faster in Aβ_42_ mice

Glutamate spillover is known to increase NMDA receptor activation, whereas glutamate uptake provides a powerful mechanism to constrain it^47–49^. Given that NMDA receptor activation in response to synaptically-released glutamate is reduced in Aβ_42_ mice (**Fig. 3**), we reasoned that this could be due to an increase in glutamate uptake capacity induced by Aβ_42_. Most glutamate transporters are expressed in astrocytes^50^. GLT-1 is the most abundant glutamate transporter in the adult brain, whereas GLAST is expressed at lower levels^51^. We hypothesized that an increase in glutamate uptake could occur in response to astrogenesis, or to a proliferation of astrocytic processes, for example. To address these concerns, we first tested whether the surface density of astrocytes in *s.r.* was changed in Aβ_42_ mice (**Fig. 5**). By using immunolabeling experiments, we showed that the density of cells immuno-positive for S100β, a protein abundantly expressed in astrocytes, was similar in WT, Sham and Aβ_42_ mice, suggesting that Aβ_42_ does not promote astrocyte proliferation 3-8 weeks post-injections (**Fig. 5A,B**). We then patched astrocytes and obtained biocytin fills, from which we derived 3D reconstructions of individual cells. The analysis showed that the astrocyte surface area and volume was similar in the three different cohorts of mice, indicating that Aβ_42_ does not induce proliferation of astrocytic processes (**Fig. 5C-F**). Patching astrocytes allowed us to analyze some key electrophysiological properties of these cells, like their resting membrane potential (**Fig. 5G,H**). On average, the resting membrane potential of astrocytes was similar in WT, Sham and Aβ_42_ mice (**Fig. 5H**). Electrical stimuli delivered to Schaffer collaterals evoked synaptically-activated transporter currents (STCs), followed by sustained potassium currents, as reported before^48,49,52^. These two components can be distinguished from one another because of their different time course and sensitivity to glutamate transporter antagonists^49^. The STC has a rapid onset and decays in tens of milliseconds; the sustained potassium current decays over a time course of seconds and its time course follows that of potassium re-equilibration in the extracellular space in response to action potential propagation^49^. The integral over time of the STC provides a loose measure for how much glutamate is taken up by astrocytes; the amplitude of the sustained potassium current varies with the number of action potentials evoked by the electrical stimulation. By plotting the relationship between these two values, we can estimate how much glutamate is taken up by astrocytes for a given number of evoked action potentials. Overall, there was no significant difference across groups (**Fig. 6I**). One thing we noticed, however, is that we could not record STCs from all patched astrocytes, especially in Aβ_42_ mice (**Fig. 5J**). In astrocytes with detectable STCs, the integral of these currents over time was similar in WT, Sham and Aβ_42_ mice, suggesting that these cells take up the same amount of glutamate, and likely express the same number of glutamate transporters (**Fig. 5K**). We used these STCs to derive temporal information about the glutamate uptake process, using a deconvolution analysis described previously^52,53^. Surprisingly, we found that glutamate clearance was faster in astrocytes from Aβ_42_ mice (**Fig. 5L,M**).

**Figure 5.**
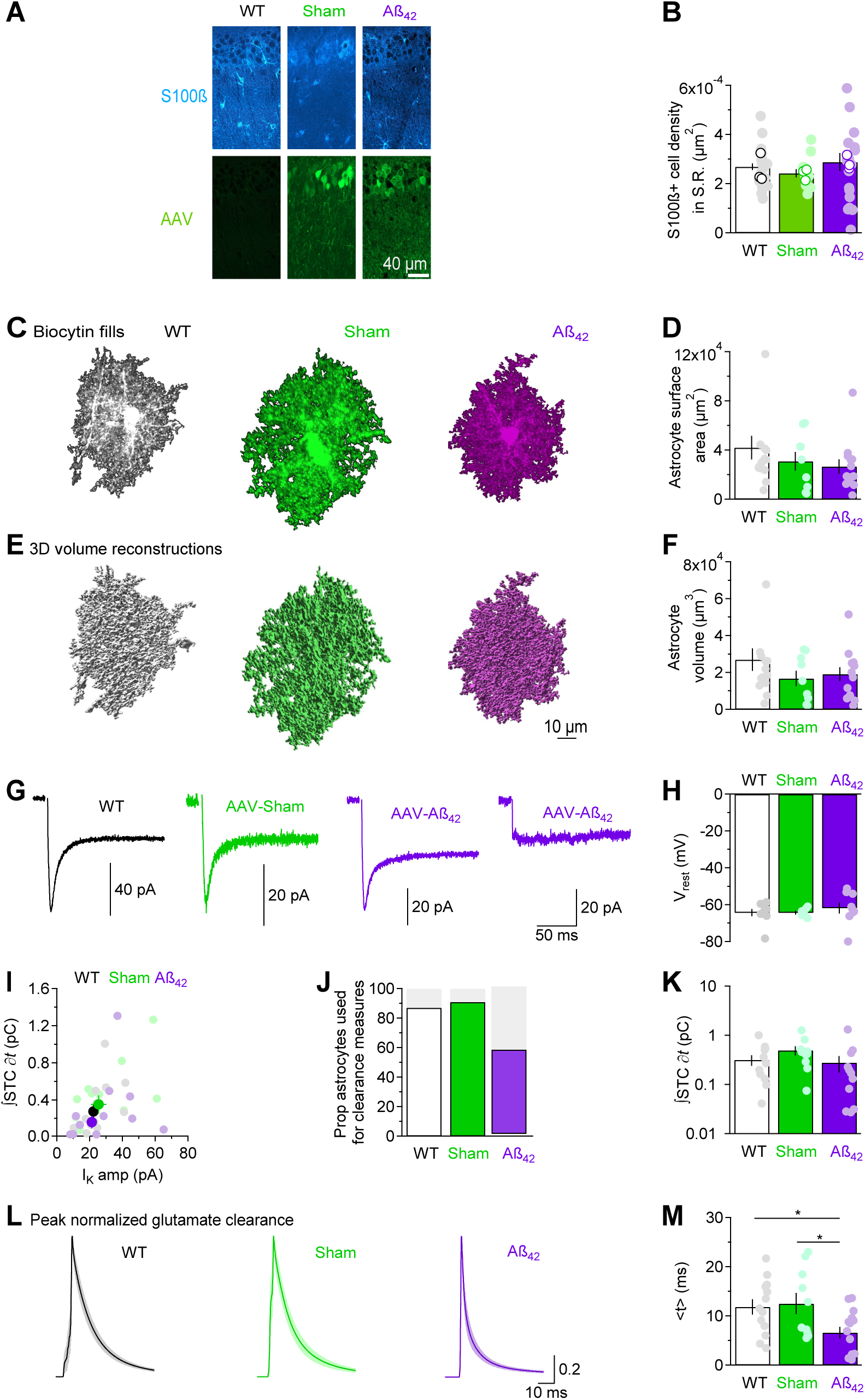
Glutamate clearance is faster in Aβ_42_ mice but can only be measured from a subset of astrocytes. (A) Immunofluorescence labeling of S100β and AAV expressing cells in hippocampal *s.r.* of WT, Sham and Aβ_42_ mice. (B) Summary of surface density of S100β expressing cells in WT (*N*=3, n=35), Sham (*N*=3, n=13) and Aβ_42_ mice (*N*=3, n=21). Empty circles refer to data collected from three different mice per group. Filled circles refer to estimates collected in individual ROIs. (C) Example of biocytin fills of astrocytes in WT, Sham and Aβ_42_ mice. (D) Summary of astrocyte surface area in WT (*N*=7, n=14), Sham (*N*=5, n=9) and Aβ_42_ mice (*N*=7, n=13). (E) Example of 3D volume reconstructions obtained from the biocytin fills using Imaris 9.2. (F) Summary of astrocyte volume measurements collected from the biocytin fills. (G) Example of electrically evoked STCs and sustained potassium currents recorded from astrocytes. In some cases, in the Aβ_42_ group, the electrical stimulation did not evoke an STC, but a potassium current could still be recorded. (H) Summary of resting membrane potential measures collected from patched astrocytes in WT (*N*=4, n=9), Sham (*N*=3, n=6) and Aβ_42_ mice (*N*=3, n=9). (I) Scatter plot describing the relationship between the integral of the STC and the amplitude of the sustained potassium current for all recordings, in WT (*N*=8, n=15), Sham (*N*=8, n=14) and Aβ_42_ mice (*N*=10, n=21). (J) Summary of the proportion of astrocytes from which we could estimate the glutamate clearance waveforms, in WT (*N*=8, n=13/15), Sham (*N*=7, n=10/14) and Aβ_42_ mice (*N*=6, n=12/21). (K) The integral over time of STCs represents the charge transfer occurring during glutamate uptake in astrocytes and is proportional to the number of glutamate molecules taken up by these cells in WT (*N*=8, n=13), Sham (*N*=7, n=10) and Aβ_42_ mice (*N*=6, n=12). (L) Example of glutamate clearance waveforms derived from WT, Sham and Aβ_42_ mice. (M) Summary of the centroid of glutamate clearance (*<T>*) measured from astrocytes in the three cohorts of mice. Data represent mean±SEM. One-way ANOVA followed by pairwise comparisons. \**p*<0.05; ** *p*<0.01; \*\*\**p*<0.001.

**Figure 6.**
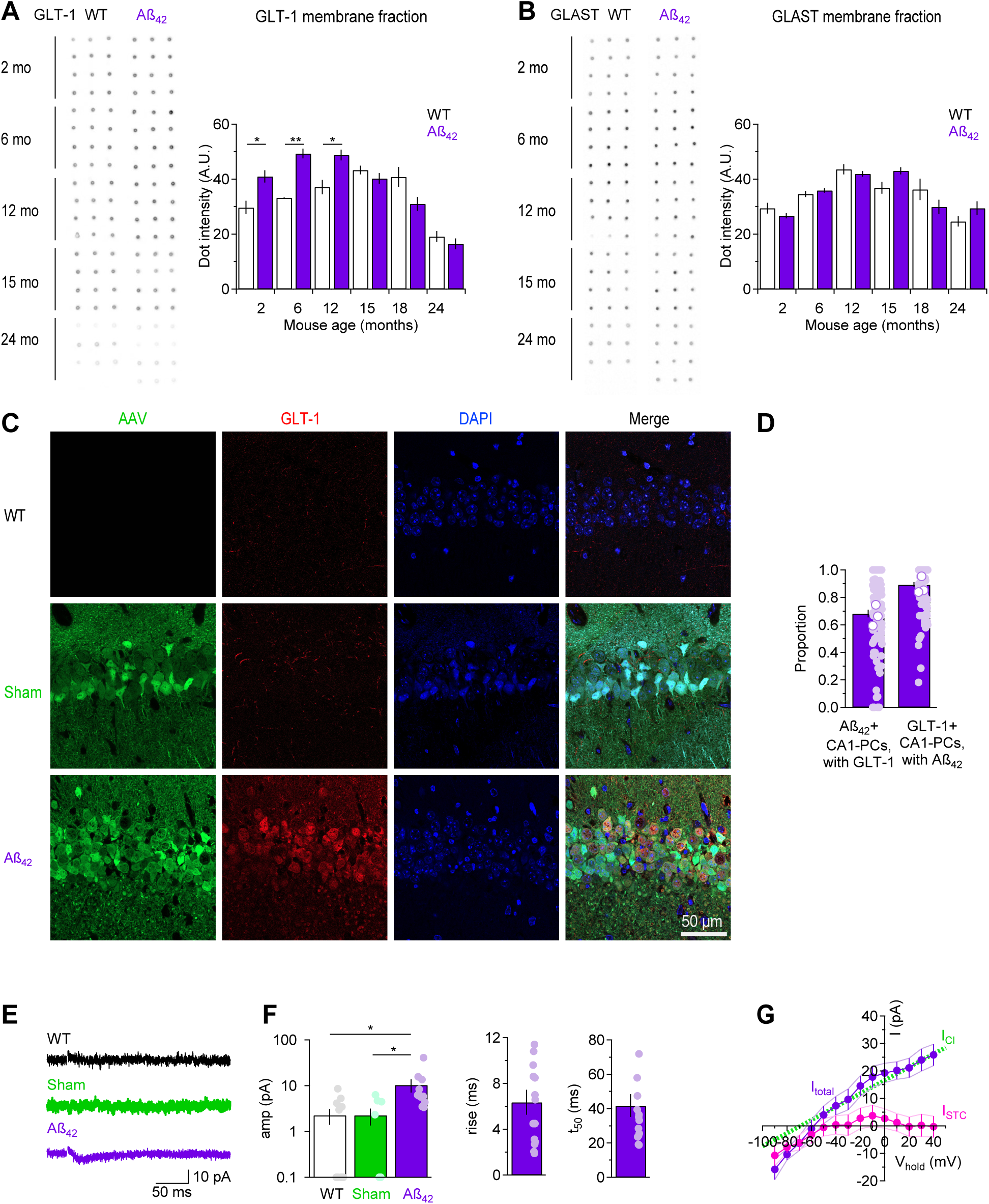
AAV-Aβ_42_ triggers ectopic GLT-1 expression in CA1-PCs. **(A)** Dot blot analysis of GLT-1 protein expression in mice aged 2-24 months, 3-4 weeks after AAV-Aβ_42_ injection. **(B)** As in A, for the glial glutamate transporter GLAST (*N*=4 for each age group). Unpaired T-test. **(C)** Immunofluorescence labeling of AAV transfection and GLT-1 expression in WT, Sham and Aβ_42_ mice. An ectopic expression of GLT-1 is only detected in Aβ_42_ mice (*N*=3, n=106). **(D)** Summary of the proportion of AAV-Aβ_42_ transfected CA1-PCs with immunoreactivity for GLT-1, and of the proportion of GLT-1 expressing CA1-PCs that are also transfected by the AAV-Aβ_42_ virus. Empty circles refer to data collected in three different mice. Filled circles refer to estimates collected in individual ROIs (*N*=3, n=106). **(E)** Representative traces showing the TFB-TBOA-sensitive currents recorded from CA1-PCs in the presence of PTX (100 µM), NBQX (20 µM) and APV (100 µM) in WT, Sham and Aβ_42_ mice. Each trace represents the average of 20 sweeps. **(F)** *Left,* Summary of the amplitude of the TFB-TBOA-sensitive current in the three cohorts of mice. *Middle,* Average 20-80% rise time of the TFB-TBOA-sensitive current in CA1-PCs of Aβ_42_ mice. *Right*, Average decay time of the TFB-TBOA-sensitive current in CA1-PCs of Aβ_42_ mice. WT (*N*=5, n=11), Sham (*N*=5, n=8), Aβ_42_ (*N*=7, n=11). One-way ANOVA followed by pairwise comparisons. **(G)** I/V profile of the TFB-TBOA-sensitive current in CA1-PCs of Aβ_42_ mice (purple). The I/V profile of the chloride current was estimated as the line that reverses at E_Cl_=-69 mV and that accounts for the entirety of the TFB-TBOA-sensitive current at V_hold_=40 mV (*green*). The putative stoichiometric in these cells was obtained by subtracting I_Cl_ from each I/V profile (*magenta*). WT (*N*=5, n=10). Data represent mean±SEM. \**p*<0.05; ** *p*<0.01; \*\*\**p*<0.001.

To determine whether this could be due to overall changes in the expression of glial glutamate transporters, we performed a dot blot analysis to measure the protein expression levels of the glial glutamate transporters GLT-1 and GLAST (**Fig. 6**). AAV-Aβ_42_ induced a significant increase in GLT-1 expression in 2 month old mice (the ones we typically used for all our experiments; **Fig. 6A**). To determine whether this effect could be age-dependent, we repeated this analysis in mice aged 6-24 months, sacrificed 3-8 weeks after the AAV-Aβ_42_ injection. AAV-Aβ_42_ lead to an increased GLT-1 expression in mice aged 2-12 months, not in older mice aged 15-24 months (**Fig. 6A**). By contrast, the expression of GLAST was similar between WT and Aβ_42_ mice aged 2-24 months (**Fig. 6B**). In both cases, the developmental trend of expression of GLT-1 and GLAST was consistent with the one we reported in previous work^51^. These findings raise an apparent paradox: on one hand the glutamate uptake capacity of astrocytes is not affected by Aβ_42_ (**Fig. 5K**), but glutamate clearance is faster (**Fig. 5L,M**) and GLT-1 expression is increased in Aβ_42_ mice (**Fig. 6A**). Could the increased GLT-1 expression in Aβ_42_ mice occur in cells other than astrocytes? Although the vast majority of GLT-1 is expressed in astrocytes, there is ultrastructural evidence that a small proportion of this protein is also expressed in neurons^54–58^. This prompted us to perform immunofluorescence experiments, to determine whether the increased expression of GLT-1 detected with dot blot experiments could be accounted for by an increased GLT-1 expression in CA1-PCs (**Fig. 6C**). Indeed, ∼68% of CA1-PCs transfected with AAV-Aβ_42_ expressed GLT-1, and ∼89% of the GLT-1-expressing CA1-PCs showed transfection with AAV-Aβ_42_ (**Fig. 6D**). If these transporters are functional in the cell soma, we should be able to record STCs from CA1-PCs of Aβ_42_ mice. To do this, we evoked EPSCs in CA1-PCs voltage-clamped at −70 mV, and blocked them with picrotoxin (100 µM), NBQX (20 µM) and APV (100 µM; **Fig. 6E,F**). In these experiments, the concentration of NBQX and APV was twice as large compared to the ones typically used in our recording conditions, to make sure the TFB-TBOA-sensitive current was not an unblocked AMPA or NMDA current. We isolated the residual currents as the currents sensitive to the broad-spectrum glutamate transporter antagonist TFB-TBOA (5 µM; **Fig. 6E,F**). These currents were significantly larger in Aβ_42_ compared to WT and Sham mice (**Fig. 6F**). Their current/voltage relationship showed an inward rectification consistent with that of a mixed non-stoichiometric Cl^-^ current, and a stoichiometric current associated with glutamate flux across the plasma membrane (**Fig. 6G**)^59,60^. Both currents require glutamate binding to the transporter. The stoichiometric current is mediated by the inward movement of 3 Na^+^, 1 H^+^ and one glutamate molecule (which carries a negative charge) and is coupled to the counter movement of 1 K^+^ or 1 Cs^+^ across the membrane^59–61^. We used the following rationale to isolate these two currents. The Cl^-^ current is zero, by definition, at the reversal potential for this ion (E_Cl_ = −69 mV). Under our experimental conditions, the electrochemical gradient for the ions co-transported with glutamate does not support a stoichiometric *outward* current. Therefore, the outward current recorded at a holding potential of +40 mV is entirely mediated by Cl^-^. These considerations allow us to estimate the I/V relationship of the TFB-TBOA-sensitive anionic current as the one described by a line connecting these two points. The stoichiometric current was then obtained by subtracting I_Cl_ from the TFB-TBOA-sensitive current recorded experimentally (**Fig. 6G**). Together, these findings suggest that Aβ_42_ promotes GLT-1 expression in CA1-PCs, increasing the rate of glutamate clearance from the extracellular space.

### Aβ_42_ increases proximal inhibition onto CA1-PCs

The data presented thus far suggest that Aβ_42_ increases the number of excitatory synaptic contacts onto CA1-PCs, but also that there is a reduced activation of NMDA receptors in Aβ_42_ mice likely due to an increased expression of GLT-1 in CA1-PCs. Whether or not this leads to an increased activity in CA1-PCs depends on other factors, including potential changes in synaptic inhibition onto CA1-PCs induced by Aβ_42_. To address this, we first recorded mIPSCs from CA1-PCs and found that their frequency was increased in CA1-PCs of Aβ_42_ mice (**Fig. 7A,B**). Their amplitude and kinetics were not altered, suggesting that Aβ_42_ does not alter the quantal size of GABAergic events (**Fig. 7B-D**). The increased mIPSC frequency is unlikely to be accounted for by an increased release probability, because the PPR of IPSCs recorded from CA1-PCs (which is inversely proportional to P_r_) was similar in WT, Sham and Aβ_42_ mice (**Fig. 7E, F**). We performed optogenetics experiments to determine whether proximal and distal inhibition were equally affected by Aβ_42_. To accomplish this, we injected a conditional AAV encoding the red shifted opsin C1V1 in hippocampal area CA1 of *Pvalb*^Cre/+^ or *Sst*^Cre/+^ mice. Some of these mice were only injected with C1V1, others also with AAV-Sham or AAV-Aβ_42_. We used the same light stimulus intensity for all experiments, to record optically-evoked IPSCs (oIPSCs) from CA1-PCs. The oIPSCs evoked by stimulating SST-INs had similar amplitude and kinetics in *Sst*^Cre/+^, *Sst*^Cre/+^-Sham and *Sst*^Cre/+^-Aβ_42_ mice (**Fig. 7G-I**). By contrast, the oIPSCs evoked by stimulating PV-INs were larger in amplitude but similar in kinetics in *Pvalb*^Cre/+^-Aβ_42_ compared to *Pvalb*^Cre/+^ and *Pvalb*^Cre/+^-Sham mice (**Fig. 7J-L**). We asked whether Aβ_42_ changed the expression of the neuronal potassium-chloride co-transporter KCC2, which is responsible for maintaining GABAergic inhibition. However, we did not detect any significant difference in the expression level of KCC2 between WT, Sham and Aβ_42_ mice (**Supp.** Fig. 3). These findings suggest that Aβ_42_ enhances proximal inhibition without altering distal inhibition onto CA1-PCs and without changing KCC2 expression.

**Figure 7.**
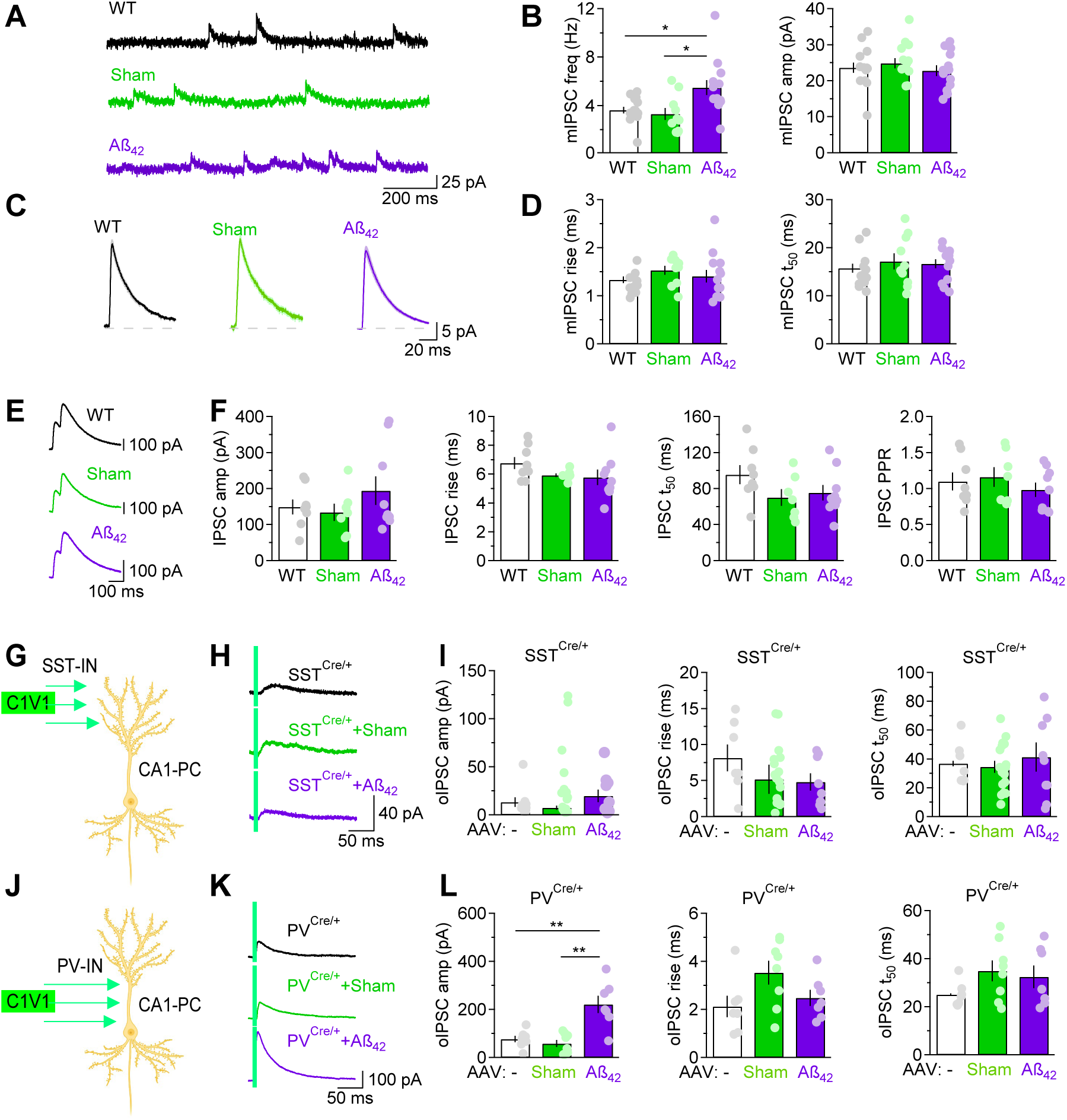
AAV-Aß_42_ leads to an early-onset increase in mIPSC frequency in CA1-PCs. **(A)** Example of mIPSC recordings in CA1-PCs of WT (*N*=4, n=15), Sham (*N*=4, n=10), and Aβ_42_ mice (*N*=3, n=13) voltage clamped at 0 mV. **(B)** Summary of mIPSC frequency (*left*) and amplitude (*right*). **(C)** Average mIPSCs recorded in CA1-PCs of WT, Sham, and Aβ_42_ mice. **(D)** As in B, for 20-80% rise time (*left*) and 50% decay time (*right*). **(E)** Average paired IPSCs recorded in CA1-PCs of WT (*N*=3, n=8), Sham (*N*=5, n=7), and Aβ_42_ mice (*N*=3, n=9). **(F)** Analysis of IPSC amplitude, 20-80% rise time, 50% decay time and PPR for evoked IPSCs. **(G)** Schematic representation of optogenetic stimulation of distal inhibition from SST-INs onto CA1-PCs. **(H)** Example of SST-IN oIPSC recorded from CA1-PCs of WT (*N*=3, n=9), Sham (*N*=6, n=17), and Aβ_42_ mice (*N*=4, n=10). The vertical cyan line represents the artifact of the optical stimulation. Each trace represents the average of 10 oIPSCs. **(I)** Summary of SST-IN mediated oIPSC amplitude (*left*), 20-80% rise time (*center*) and 50% decay time (*right*). **(J-L)** As in (G-I), for oIPSCs evoked by optogenetic stimulation of PV-INs (WT *N*=3, n=7; Sham *N*=3, n=8, Aβ_42_ *N*=3, n=7). Data represent mean±SEM. One-way ANOVA followed by pairwise comparisons. \**p*<0.05; ** *p*<0.01; \*\*\**p*<0.001.

To determine how these effects shaped the temporal sequence of excitation-inhibition (E/I) in hippocampal area CA1, we performed a series of experiments using 64 electrode multi-electrode arrays (MEAs). One of the MEA electrodes was used to deliver current stimuli to Schaffer collaterals (1 mA×80 µs; **Fig. 8A**). This evoked a fiber volley (FV), followed by an E/I sequence that differed between WT/Sham and Aβ_42_ mice (**Fig. 8B**). Specifically, the evoked field EPSP (fEPSP) was larger and shorter-lived in Aβ_42_ mice (**Fig. 8B**). Accordingly: *(i)* the ratio between the fEPSP slope and the fiber volley amplitude, which provides a rough estimate of Schaffer collateral activation and evoked release, was larger in Aβ_42_ than in WT and Sham mice (**Fig. 8C, left**) and *(ii)* the centroid of the fEPSP (i.e., its center of mass) was shorter in Aβ_42_ than in WT and Sham mice (**Fig. 8C, right**). The analysis of the temporal progression of the propagation of the extracellular field confirmed that this decayed more rapidly in Aβ_42_ mice (**Fig. 8D-E**). Overall, the charge transfer of the fEPSP was similar across the three mouse groups (**Fig. 8F, left**), because the larger fEPSPs in Aβ_42_ mice were curtailed by larger fIPSPs (**Fig. 8F, right**).

**Figure 8.**
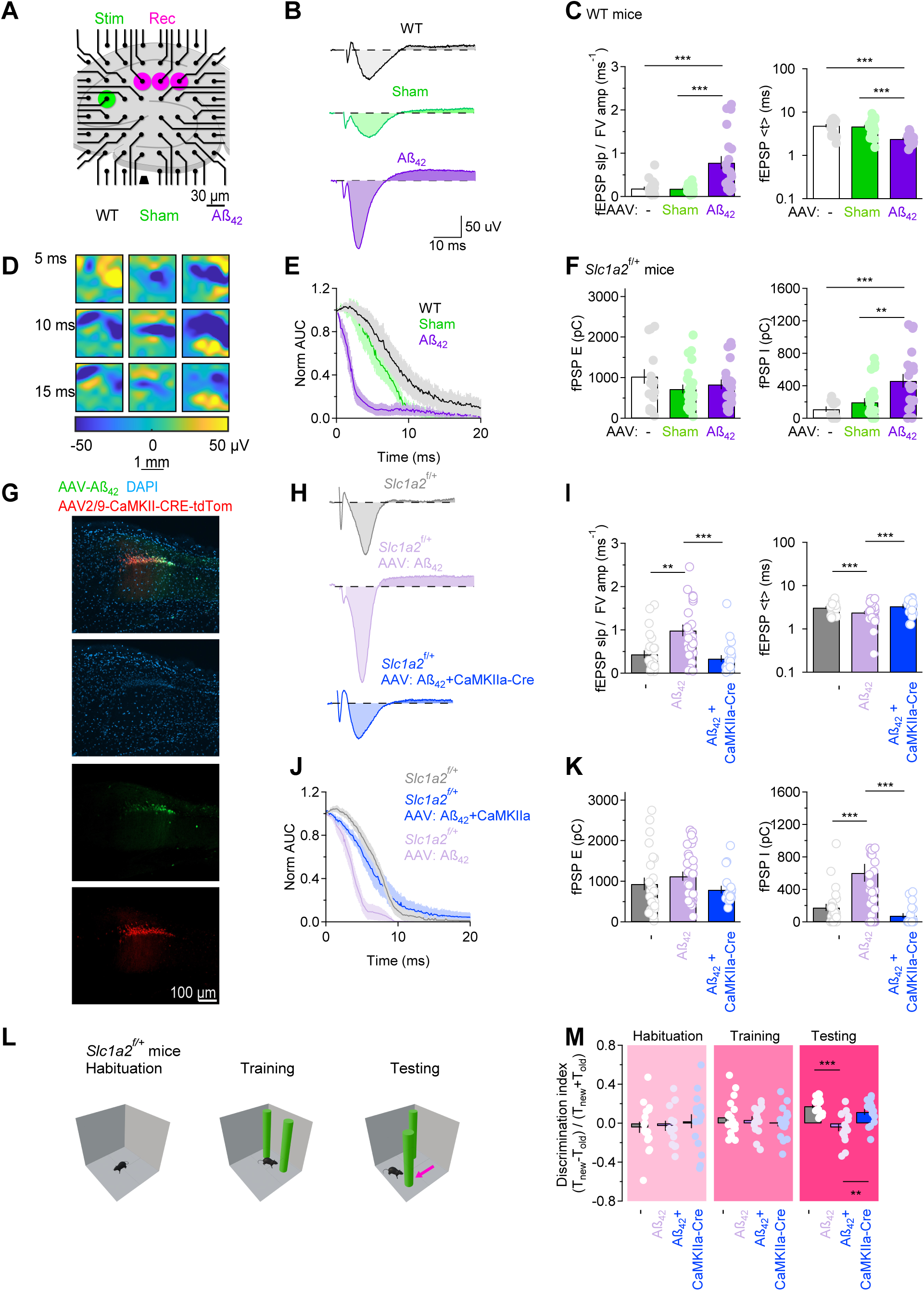
AAV-Aß_42_ leads to larger but shorter-lived excitation in hippocampal area CA1. **(A)** Schematic representation of MEA recording system, with stimulating and selected recording electrodes highlighted in green and magenta, respectively. The ground electrode is represented as a black trapezoid. **(B)** Representative field recordings obtained from one of the selected recording electrodes for slices from WT, Sham and Aβ_42_ mice. Each trace represents the average of 30 consecutive trials, following TTX background subtraction. **(C)** *Left*, Summary of the fEPSP slope and fiber volley ratio measured in slices from WT (*N*=3, n=18), Sham (*N*=4, n=20), and Aβ_42_ mice (*N*=3, n=24). *Right*, Summary of the fEPSP centroid (i.e., the center of mass) calculated in the three cohorts of mice. **(D)** Snapshots of MEA extracellular field potential propagation in a hippocampal slice of WT, Sham and Aβ_42_ mice. The snapshots were collected 5, 10 and 15 ms after the stimulus artifact. The slice orientation was as shown in panel A. The dark blue area identifies the regions with the largest downward fEPSP. **(E)** Decay of the peak normalized fEPSP amplitude in the three mouse cohorts (WT n=7; Sham n=9, Aβ_42_ n=7). **(F)** *Left*, Charge transfer of the fEPSPs, corresponding to the light shaded areas in panel B. *Right*, Charge transfer of the fIPSPs, corresponding to the dark shaded areas in panel B**. (G)** Immunofluorescence labeling of hippocampal sections from *Slc1a2^f/+^* mice injected with AAV-Aβ_42_ (*green*) and AAV-CaMKII-CRE-tdTom (*red*). **(H)** Representative field recordings obtained from one of the selected recording electrodes for slices from *Slc1a2^f/+^*mice (gray), *Slc1a2^f/+^*mice injected with AAV-Aβ_42_ (*light purple*) and *Slc1a2^f/+^* mice injected with AAV-Aβ_42_ and AAV-CaMKII-CRE-tdTom (*blue*). Each trace represents the average of 30 consecutive trials, following TTX background subtraction. **(I)** *Left*, Summary of the fEPSP slope and fiber volley ratio measured in slices from *Slc1a2^f/+^* mice (*N*=5, n=25), *Slc1a2^f/+^* mice injected with AAV-Aβ_42_ (*N*=5, n=31), and *Slc1a2^f/+^*mice injected with AAV-Aβ_42_ and AAV-CaMKII-CRE-tdTom (*N*=3, n=21). *Right*, Summary of the fEPSP centroid (i.e., the center of mass) calculated in the three cohorts of mice. **(J)** Decay of the peak normalized fEPSP amplitude in the three mouse cohorts (*Slc1a2^f/+^* n=8; *Slc1a2^f/+^* mice injected with AAV-Aβ_42_ n=6, *Slc1a2^f/+^*mice injected with AAV-Aβ_42_ and AAV-CaMKII-CRE-tdTom n=8). **(K)** *Left*, Charge transfer of the fEPSPs, corresponding to the light shaded areas in panel H. *Right*, Charge transfer of the fIPSPs, corresponding to the dark shaded areas in panel H**. (L)** Summary of object location test, showing that *Slc1a2^f/+^* mice injected with Aβ_42_ do not discriminate the location of a displaced object (*Slc1a2^f/+^* N=9, n=20; *Slc1a2^f/+^*mice injected with AAV-Aβ_42_ *N*=7, n=15, *Slc1a2^f/+^* mice injected with AAV-Aβ_42_ and AAV-CaMKII-CRE-tdTom N=7, n=16). Data represent mean±SEM. One-way ANOVA followed by pairwise comparisons. \**p*<0.05; ** *p*<0.01; \*\*\**p*<0.001.

A fundamental unknown in these experiments is whether these changes in activity propagation are directly triggered by Aβ_42_ or rather by the Aβ_42_-induced increase in GLT-1 expression in CA1-PCs. To test this hypothesis, we performed MEA and behavioral experiments on *Slc1a2*^f/+^ mice. In these mice, targeted Cre-LoxP recombination can be used to reduce GLT-1 expression in different cell types. We collected data from: *(i)* non-injected *Slc1a2*^f/+^ mice; *(ii) Slc1a2*^f/+^ mice injected with AAV-Aβ_42_; *(iii) Slc1a2*^f/+^ mice injected with both AAV-Aβ_42_ and AAV-CaMKIIa^Cre^ (**Fig. 8G**). The results showed that Aβ_42_ altered hippocampal activity propagation in *Slc1a2*^f/+^ mice as it did in WT mice (**Fig. 8H-K**). However, Aβ_42_ no longer altered the network activity in area CA1 if AAV-Aβ_42_ was injected in mice lacking neuronal GLT-1 (**Fig. 8H-K**). To determine whether spatial memory could be rescued in AAV-Aβ_42_ injected mice that did not express neuronal GLT-1, we repeated the novel object location test in *(i)* non-injected *Slc1a2*^f/+^ mice; *(ii) Slc1a2*^f/+^ mice injected with AAV-Aβ_42_; *(iii) Slc1a2*^f/+^ mice injected with both AAV-Aβ_42_ and AAV-CaMKIIa^Cre^ (**Fig. 8L,M**). In these experiments, we found that the only mouse group showing a memory loss was *Slc1a2*^f/+^ mice injected with AAV-Aβ_42_. Together, these findings indicate that the changes in the population activity of hippocampal area CA1 and the cognitive disruption induced by AAV-Aβ_42_ are largely mediated by an increase in GLT-1 expression in CA1-PCs.

## DISCUSSION

Although there is substantial evidence that Aβ disrupts both excitatory and inhibitory synapses in AD pathology, our understanding of how specific Aβ peptides contribute to synaptic and cellular dysfunction in the early stages of AD remains limited^62–65^. In this work, we addressed this concern by taking advantage of an AAV vector-based approach, which we used to promote Aβ_42_ expression in hippocampal area CA1. Our findings indicate that Aβ_42_ induces early changes in hippocampal synaptic activity, which include an increases the number of excitatory synaptic contacts onto CA1-PCs and a reduced activation of NMDA versus AMPA receptors at Schaffer collateral synapses. Aβ_42_ increases proximal inhibition onto CA1-PCs, while also promoting the expression of the glutamate transporter GLT-1 in these cells. Together, these effects contribute to alter the excitability of hippocampal circuits and disrupt spatial memory. Notably, these early effects of Aβ_42_ appear to be largely mediated by an Aβ_42_-induced increase in the expression of the glutamate transporter GLT-1 in CA1-PCs.

### Prior and current data

Our finding of an *increased* GLT-1 expression induced by AAV-Aβ_42_ may seem at odds with others showing *reduced* GLT-1 expression and uptake in the cortex and hippocampus of AD patients^66–68^. Our own previous *in vitro* experiments, in which we acutely applied a high dose of Aβ_42_ (0.5 µM) to hippocampal slices, showed a *reduction* in GLT-1 expression and a slower rate of glutamate uptake from astrocytes^69^. There are numerous technical differences between past and present works. *First*, the concentration of soluble Aβ_42_ in our ELISA samples is ∼50 pg/µg protein (**Fig. 1D**). If the brain tissue contains ∼97.8 gm/kg protein^70^ and has a density of 1.05 gm/cm^3^ ^71^, the concentration of soluble Aβ_42_ in the RIPA-treated samples is ∼1 µM. Although these numbers are in the same ballpark, we expect only part of the Aβ_42_ in the ELISA tests to derive from the extracellular pool (the rest being intracellular). For this and other reasons listed below, it is challenging to make a reliable close comparison between acute and AAV work. *Second*, acute slice applications does not lead to significant Aβ_42_ aggregation, whereas most Aβ_42_ expressed via the AAV approach tends to form aggregates, so we are likely looking at the effect of different Aβ_42_ aggregates on glutamate uptake (**Fig. 1D**). *Third,* acute applications are brief (20-40 min), whereas the AAV mimics more of a chronic state. *Fourth,* the AAV approach allows us to focus on an earlier time point in the progression of the Aβ pathology compared to the time point that can be analyzed post-mortem in humans. For all these reasons, the most parsimonious interpretations of these different datasets is that that should be considered as such.

### Age-dependent effects of Aβ_42_

Interestingly, in our experiments, Aβ_42_ does not induce upregulation of GLT-1 in mice older than one year (**Fig. 6A**). We do not know the exact signaling mechanisms mediating this upregulation, but we hypothesize that the expression of some of their key components may decline with age, potentially making the hippocampus refractory to the deleterious effects of Aβ_42_. In this scenario, AD would emerge as an early-onset disease, with clinical manifestations that are significantly delayed compared to its latent cellular progression.

### What is neuronal GLT-1 good for in the healthy and in the AD brain?

The finding that Aβ_42_ induces GLT-1 expression in neurons is particularly noteworthy, in our opinion. In the adult and healthy brain, glutamate transporters are abundantly expressed in astrocytes. However, GLT-1 mRNA has also been detected in neurons of the rat hippocampus (particularly CA3-PCs), striatum somatosensory cortex and human cortex^54,56–58,72–77^. Even though only 5-6% of all hippocampal GLT-1 protein is in neurons^54,78^, immunoreactivity for GLT-1 can be detected in 14-29% of excitatory axon terminals^54^ (this proportion is much higher in synaptosomes^78,79^). This small amount of neuronal GLT-1 mediates more than half of the uptake of the exogenous substrate D-Asp in slices^55,78,79^. This raises the possibility that the functional properties of neuronal GLT-1 might be different from those of astrocytic GLT-1, perhaps due to post-translational modifications that render it more active than astrocytic GLT-1^55,80^. If this were true, we would expect neuronal GLT-1 to have an important functional relevance for regulating hippocampal-dependent phenotypes. In recent times, the question of the function of neuronal GLT-1 was explored using conditional knockout mice, where GLT-1 expression in neurons was prevented through Cre expression driven by the synapsin I promoter^78,81^. These mice had normal survival, weight gain, and no seizures, but displayed alterations in synaptic mitochondrial metabolism, late-onset spatial reference long-term memory deficits, suggesting that indeed neuronal GLT-1 exerts an important physiological role^78,79,81^. It was suggested that neuronal GLT-1 may be involved in the regulation of vulnerability to excitotoxicity and altered glucose metabolism via mitochondrial superoxide production^82,83^. Based on our data, we would like to suggest that the small proportion of neuronal GLT-1 might serve as a reservoir pool of protein that can be upregulated by peptides that accumulate in the AD brain, like Aβ_42_. This could represent an important compensatory mechanism to counteract the increased glutamatergic drive onto CA1-PCs, limiting NMDA receptor activation and excitotoxicity. Consistent with this hypothesis, recent slice physiology data show that Aβ-induced impairment of hippocampal long-term plasticity is suppressed by neuronal GLT-1 knockout, perhaps due to its ability to supply glutamate to synaptic mitochondria^79,83,84^.

From an electrophysiological standpoint, glutamate transporter currents in CA1-PCs have never been recorded, but this finding should be interpreted with caution. Being unable to record a transporter current from the soma does not rule out the possibility that these currents may occur in dendrites and be electrotonically filtered as they propagate towards the soma. This phenomenon has been described extensively for the neuronal glutamate transporter EAAC1 and could also apply to neuronal GLT-1^48,85–87^. Like other glutamate transporter isoforms and homologs from archaebacteria to mammals, GLT-1 exhibits a dual function: it acts as a secondary active glutamate transporter and as a glutamate-gated anion channel permeable to Cl^-^. Given that GLT-1 contributes to Cl^-^ homeostasis under physiological conditions, it is possible that an Aβ_42_-induced increase in GLT-1 expression in CA1-PCs may contribute to regulate Cl^-^homeostasis in these cells^88–90^. In fact, there is clinical evidence showing that: *(i)* the expression of the K^+^/Cl^-^ cotransporter KCC2, involved with Cl^-^ homeostasis, is reduced in the frontal lobe of patients with idiopathic AD and in hippocampal area CA1 of 5xFAD and *App*^NL-G-F^ mice; *(ii)* restoring neuronal Cl^-^ extrusion via KCC2 in mice reverses cognitive decline and protects against cortical hyperactivity^91,92^. In our hands, Aβ_42_ does not induce any detectable change in KCC2 expression (**Supp.** Fig. 3). If neuronal GLT-1 could serve to regulate Cl^-^ homeostasis, it would represent an important promising molecular target to restore synaptic and neuronal function in AD pathology. Interestingly, in heterologous expression systems, GLT-1 exhibits a small Cl^-^ conductance^59^. Since the Cl⁻ permeability of glutamate transporters is not strictly determined by their genetic sequence and can be induced by small structural changes in the protein, further studies are needed to clarify how these transporters contribute to Cl⁻ homeostasis in native environments^93,94^.

### Inhibition in AD

In AD patients, changes in the subunit composition of GABA_A_ receptors has been observed^95–97^. Although in APP overexpressing mice, with APP driven by strong non-specific promoters, there is a significant loss of PV- and SST-INs^98–101^, this is not detected in *App*^NL-F^ mice^102^. Here, PV- and SST-INs are not lost, but PV synapses targeting the axon initial segment are significantly larger^102^. Recent findings suggest however that the landscape of IN-specific changes induced by APP may be more complex than previously thought, with a regional and cell-type-specific susceptibility to the progression of AD pathology that deserves further investigation. In our AAV-Aβ_42_ model, we detected changes in PV- but not SST-inhibition onto CA1-PCs. Because PV-INs can tightly regulate the firing output of CA1-PCs and the stability and precision of place fields^103^, while SST-INs provide instructive signals for CA1-PCs to assess the saliency of memory information propagated through the hippocampal circuit^104^, compromising the function of these cells could provide specific cues to decode behavioral abnormalities in prodromal AD.

### Neuronal hyperactivity as a key prelude to neuronal loss

Neuronal hyperactivity, and in some cases seizures, occur in pre-symptomatic AD patients and in mice with AD-related mutations. This can exacerbate AD pathology, decrease KCC2 expression and promote Aβ_42_ accumulation, before progressing to hypoactivity in later stages of the disease^6,105^. At the cellular level, hyperactivity is pronounced in neurons located near Aβ plaques (<60 µm), and its onset is associated with the onset of cognitive decline^4,63^. Because its timing precedes that of neuronal loss, its prevention provides a critical window for early detection and intervention for AD. Encouraging data show that reversing hyperactivity in brain regions affected by early Aβ deposition reduces Aβ accumulation in the same region and prevents the spread of Aβ pathology to other regions of the brain^106,107^. This hyperactivity has been proposed to be contributed by multiple factors, including disrupted Ca^2+^ homeostasis, changes in glutamatergic transmission, Aβ and phopsho-tau deposition, astrocytes and interneurons dysfunction. Our findings support the key role of Aβ_42_ in this process.

## RESOURCE AVAILABILITY

All data analyzed for this paper have been deposited into an Open Science Framework repository (https://osf.io/b48my/). Original data and custom scripts used for this study are available from the lead contact upon request.

## ACKNOWLEDGEMENTS

The authors would like to thank Dr. Phillip J. Albrecht for managing and genotyping the mouse colony, and Dr. Janet L. Paluh and Maria B. Paredes-Espinosa for sharing their MEA equipment with us. Mice aged 18 months and older were a generous gift from the NIH Rodent Ordering System for NIA awardees. This work was supported by the NIH grant R01AG075338 to P.A.R., M.D., D.G.C., A.S; NIH grant R03AG070766 to

P.A.R. and P30HD018655; NSF grant IOS2011998 to A.S. Confocal images were acquired using a Zeiss LSM980 system with Airyscan 2 funded by the NIH SIG grant S10OD028600.

## AUTHOR CONTRIBUTIONS

Conceptualization: A.S. Methodology: M.D., A.S. Software: A.S.

Validation: A.S.

Formal analysis: M.A.P., M.D., A.S.

Investigation: P.H.W., T.J.R., M.A.P., E.D.C., A.K.M., L.F.R., G.C.T., N.A., S.A., H.M., I.L.T, U.H., B.R.T., A.S.

Resources: P.A.R., M.D., A.S. Data curation: A.S.

Writing – Original draft: A.S.

Writing – Review and editing: P.A.R., A.S. Visualization: A.S.

Supervision: A.S.

Project Administration: A.S.

Funding acquisition: P.A.R., M.D., D.G.C., A.S.

## SUPPLEMENTARY FIGURE LEGENDS

**Supplementary** Figure 1**. Aβ_42_ does not induce cell death 3-8 weeks after injection.** The panel shows representative images from WT mice hippocampal slices maintained in hypoxia conditions (i.e., no bubbling with 95% O_2_ / 5% CO_2_) for 30 min (first column) and in carbogenated conditions (second column). Slices from Sham (third column) and Aβ_42_ mice (fourth column) were also kept under constant 95% O_2_ / 5% CO_2_ perfusion. Color merged images are shown in the top row. AAV (second row), caspase-3 (third row), and DAPI labeling (fourth row) are shown for all samples.

**Supplementary** Figure 2**. AAV-Aß_42_ does not change the dendritic arborization of CA1-PCs. (A)** Example of biocytin fills of CA1-PCs in WT (*left*), Sham (*green*) and Aβ_42_ mice (*purple*). **(B)** 2D projections of 3D morphological reconstructions of CA1-PCs in the same three cohorts of mice. **(C)** Sholl analysis of CA1-PCs, showing the number of intersections formed by CA1-PCs at increasing distances from the soma, in WT (*N*=10, n=17), Sham (*N*=13, n=18) and Aβ_42_ mice (*N*=10, n=15). **(D)** Cumulative distributions for the Sholl analysis shown in C. **(E)** Summary of the total number of intersections formed by the dendritic branches of biocytin-filled CA1-PCs in WT, Sham, and Aβ_42_ mice. **(F)** Average number of intersections formed by CA1-PC dendrites in the same three groups of mice. **(G)** Summary of the maximum distance of dendrites from the soma of CA1-PCs. Data represent mean±SEM. One-way ANOVA followed by pairwise comparisons. \**p*<0.05; ** *p*<0.01; \*\*\**p*<0.001.

**Supplementary** Figure 3**. Aβ_42_ does not change KCC2 expression. (A)** Immunolabeling experiments to detect KCC2 expression (*red*) in WT, Sham and Aβ_42_ mice. The yellow arrowheads point to transfected CA1-PCs. **(B)** Line profiles collected across the plasma membrane of AAV-transfected CA1-PCs, in WT (*N*=6, n=260), Sham (*N*=5, n=253) and Aβ_42_ mice(*N*=5, n=250). Data represent mean±SEM. \**p*<0.05; ** *p*<0.01; \*\*\**p*<0.001.

## KEY RESOURCES TABLE

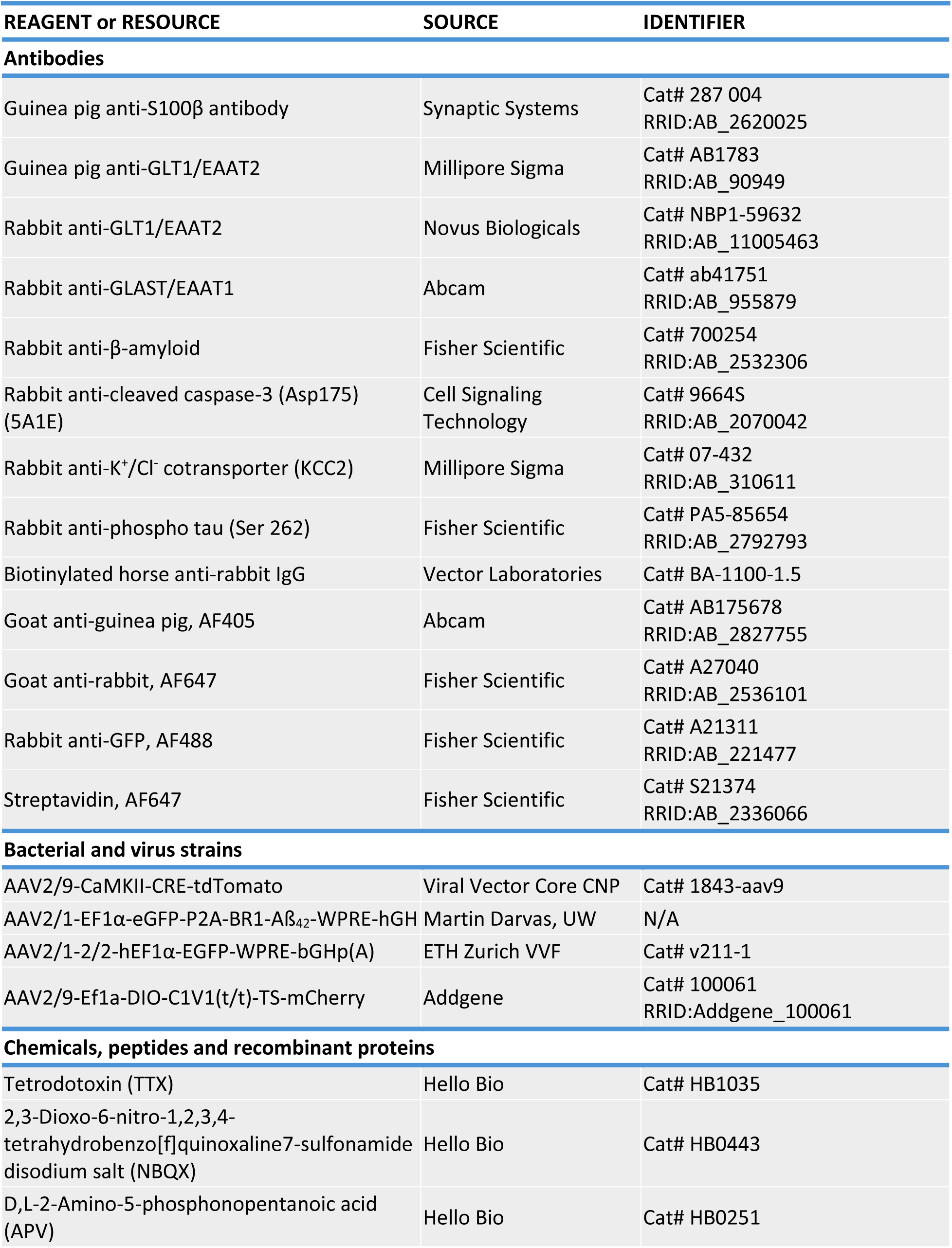
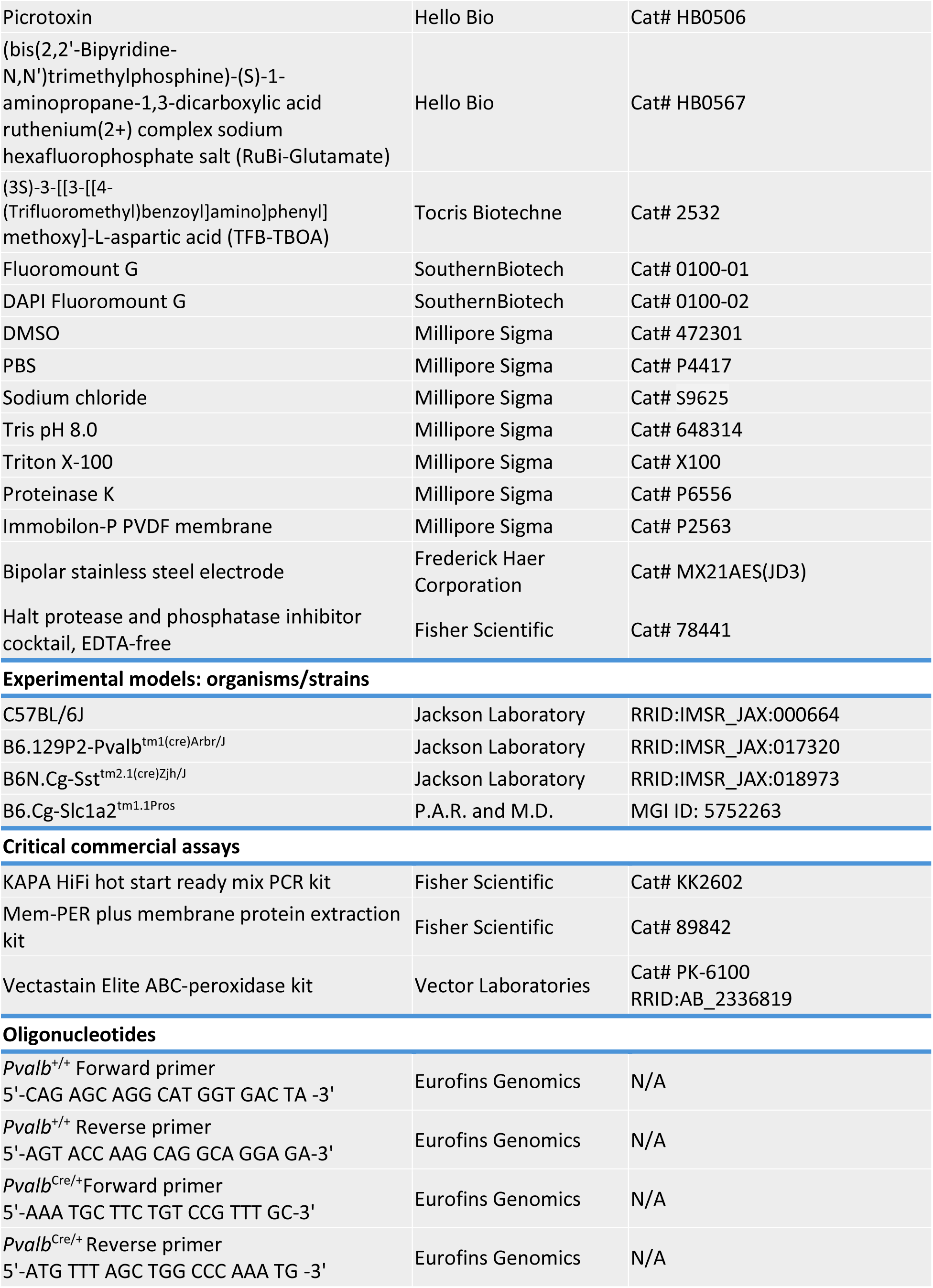
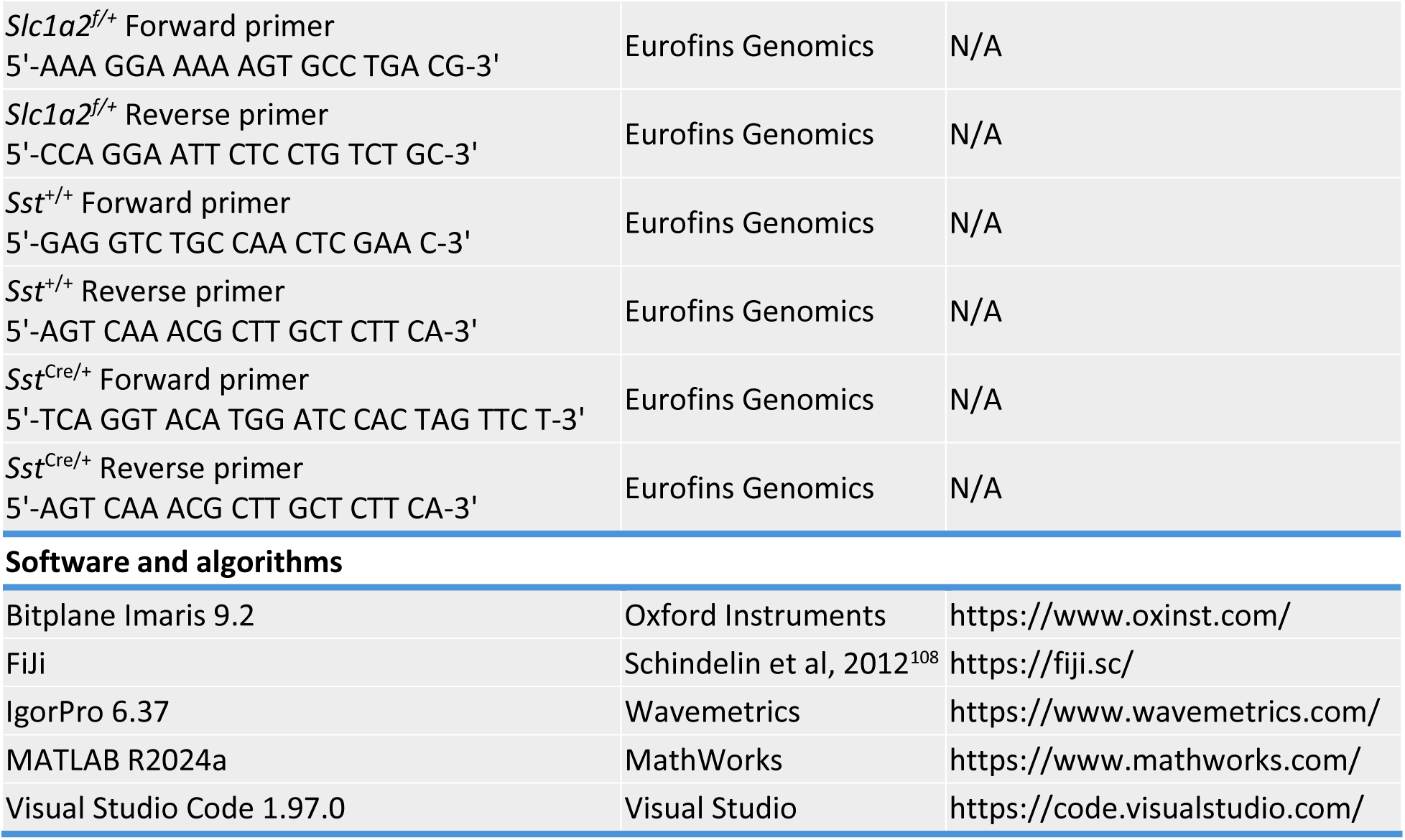

### STAR ★ METHODS

#### Mice

The **Key Resource Table** lists the mouse lines used in this study. All mice (*Mus musculus*), males and females, were group housed and kept under a 12 hr light cycle (7:00 AM on, 7:00 PM off) with food and water available *ad libitum*. C57BL/6J (WT), B6.129P2-*Pvalb*^tm1(cre)Arbr/J^ (PV^Cre/+^) and B6N.Cg-*Sst*^tm2.1(cre)Zjh/J^ (SST^Cre/+^) mice were purchased from the Jackson Laboratory (Bar Harbor, ME) and bred in house. B6.Cg-*Slc1a2*^tm1.1Pros^ mice were obtained from P.A.R. and M.D^78^. Slc1a2^f/+^ mice were obtained by crossing heterozygous with WT mice. Genotyping was performed on toe or tail tissue samples of P7-10 mice. Briefly, tissue samples were digested at 55°C overnight with shaking at 330 rpm in a lysis buffer containing the following (in mM): 100 Tris base pH 8, 5 EDTA, 200 NaCl, 0.2% SDS and 50 µg/ml proteinase K. Following heat inactivation of proteinase K at 97°C for 10 min, samples were centrifuged at 13,000 rpm for 10 min at 4°C. Supernatant DNA samples were diluted 1:1 with nuclease-free water. The PCR primers used for PV^Cre/+^ and SST^Cre/+^ genotyping were purchased from Eurofins Genomics (Louisville, KY), and their nucleotide sequences are listed in the **Key Resource Table**. PCR was carried out using the KAPA HiFi Hot Start Ready Mix PCR Kit (KAPA Biosystems, Western Cape, South Africa). Briefly, 12.5 µl of 2× KAPA HiFi Hot Start Ready Mix was added to 11.5 µl of a diluted primer mix (0.5-0.75 µM final for each primer) and 1 µl of diluted sample DNA. The PCR cycling protocol for all mutants is described in **Table 1**.

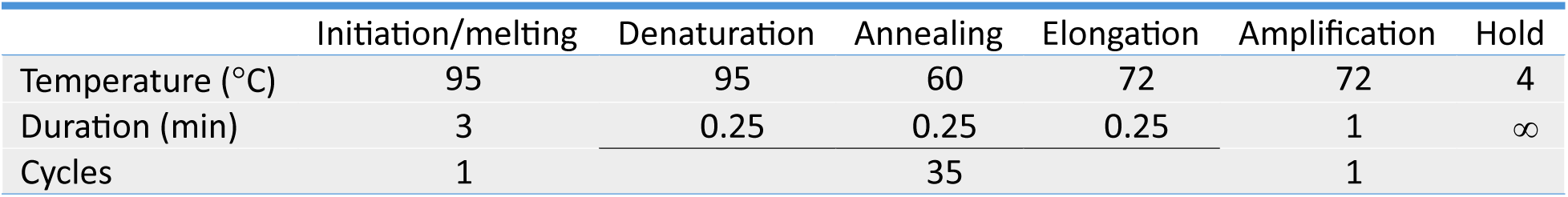

|  | Initiation/melting | Denaturation | Annealing | Elongation | Amplification | Hold |
| --- | --- | --- | --- | --- | --- | --- |
| Temperature (°C) | 95 | 95 | 60 | 72 | 72 | 4 |
| Duration (min) | 3 | 0.25 | 0.25 | 0.25 | 1 | ∞ |
| Cycles | 1 |  | 35 |  | 1 |  |

#### Stereotaxic intracranial injections and optogenetics

Each AAV was injected into hippocampal area CA1 of WT mice of either sex aged P14-16. We delivered 200 nl in each hemisphere of AAV-Sham (diluted 1:10), AAV-Aβ_42_ (undiluted), AAV2/9-Ef1a-DIO-C1V1(t/t)-TS-mCherry (undiluted) or AAV-CaMKII-Cre-TdTomato (undiluted). For the stereotaxic injections, mice were anesthetized with isoflurane (induction: 5% in 100% O_2_ at 1-2 l/min; maintenance: 3% in 100% O_2_ at 1-2 l/min) and placed in the stereotaxic frame of a motorized drill and injection robot (Neurostar GmbH, Tübingen, Germany). After making a skin incision and thinning the skull under aseptic conditions, we injected the viral constructs bilaterally in hippocampal area CA1 using a Hamilton syringe at a rate of 50 nl/min. The injection coordinates from bregma of area CA1 were AP: −1.9 mm, ML:±1.6 mm, DV: 1.4 mm. After the stereotaxic injections, the mice were returned to their home cage and used for slice physiology experiments 3-6 weeks after surgery.

#### Acute slice preparation

Acute parasagittal slices of the mouse brain were obtained from WT, PV^Cre/+^ and SST^Cre/+^ mice of either sex (5-8 week old), deeply anesthetized with halothane and decapitated in accordance with SUNY Albany Animal Care and Use Committee guidelines. The mice were transcardially perfused with 10 ml of ice-cold slicing solution bubbled with 95% O_2_/5% CO_2_ containing the following (in mM): 93 N-Methyl-D-glucamine (NMDG), 2.5 KCl, 1.2 NaH_2_PO_4_, 30 NaHCO_3_, 20 N-2-hydroxyethylpiperazine-N-2-ethane sulfonic acid (HEPES), 25 glucose, 5 L-ascorbic acid, 3 myo-inositol, 3 sodium pyruvate, 10 MgCl_2_, 0.5 CaCl_2_, 320 mOsm, pH7.4. Transverse slices of the mouse hippocampus (250 µm thick) were prepared using a vibrating blade microtome (VT1200S; Leica Microsystems, Buffalo Grove, IL). Once prepared, the slices were stored in the NMDG-based solution in a submersion chamber at 36°C for 10 min. They were then transferred to a storage solution bubbled with 95% O_2_/5% CO_2_ containing (in mM): 119 NaCl, 2.5 KCl, 1 NaH_2_PO_4_, 1.3 MgSO_4_·7H_2_O, 4 MgCl_2_, 26.2 NaHCO_3_, 0.5 CaCl_2_, 20 glucose, 320 mOsm, pH7.4. The slices were maintained in this storage solution at RT for at least 30 min and up to 5 hr.

#### Patch-clamp electrophysiology, optogenetics and uncaging experiments

Unless otherwise stated, the recording solution contained the following (in mM): 119 NaCl, 2.5 KCl, 1.2 CaCl_2_, 1 MgCl_2_, 26.2 NaHCO_3_, and 1 NaH_2_PO_4_, 22 glucose, 300 mOsm, pH7.4. Hippocampal area CA1 was identified under bright field illumination using an upright fixed-stage microscope (BX51 WI; Olympus, Center Valley, PA). Neurons and astrocytes were identified under infrared-differential interference contrast. When recording electrically evoked IPSCs, the stimulating and recording electrodes were both placed in *s.r.* ∼100 µm away from each other. Electrical stimulation was obtained by delivering constant voltage electrical pulses (50 µs) through a stimulating bipolar stainless-steel electrode (Cat# MX21AES(JD3); Frederick Haer Corporation, Bowdoin, ME). Electrical stimuli were applied every 10 s or 20 s, when recording AMPA and NMDA EPSCs, respectively. Optically evoked IPSCs were evoked using 5 ms-long light pulses generated by a SOLA-SE light engine (Lumencor, Beaverton, OR), filtered using a TRITC filter set (542/570/620 nm). The light power at the sample plane was ∼250 µW. The light pulses were delivered at intervals of 30 s using whole-field illumination through a 40× water immersion objective (LUMPLFLN40XW; Olympus, Center Valley, PA). Flash photolysis experiments were performed in the continued presence of picrotoxin (100 µM). RuBi-Glutamate (50 µM) was perfused in the recording chamber and uncaged using the output of a light engine (Lumencor, Beaverton, OR), filtered with a FITC filter set (469/497/525 nm). Each light pulse was 5 ms long, was delivered every 15 s, and had ∼500 µW at the sample plane. We used the same light stimuli to test slices from naïve WT, AAV-Sham and AAV-Aβ_42_ injected mice. To boost mEPSC and mIPSC frequency, we raised the extracellular CaCl_2_ concentration to 4 mM. For astrocyte recordings, AMPA, NMDA and GABA_A_ receptors were blocked with NBQX (10 µM), D,L-APV (50 µM) and pictrotoxin (100 µM), respectively. Whole-cell recordings from CA1-PCs and astrocytes were made with patch pipettes containing (in mM): 120 CsCH_3_SO_3_, 10 EGTA, 20 HEPES, 2 MgATP, 0.2 NaGTP, 5 QX-314Br, 290 mOsm, pH 7.2. For all electrophysiology experiments, the resistance of the recording electrode was 5-7 MOhm and was monitored throughout the experiments. Data were discarded if the resistance changed >20% during the experiment. When recording excitatory currents, picrotoxin (100 µM) was added to the recording solution to block GABA_A_ receptors. To isolate inhibitory currents, AMPA and NMDA receptors were blocked with NBQX (10 µM) and D,L-APV (50 µM), respectively. All recordings were obtained using a Multiclamp 700B amplifier and filtered at 5 KHz (Molecular Devices, San Jose, CA), converted with an 18-bit 200 kHz A/D board (HEKA, Holliston, MA), digitized at 10 KHz, and analyzed offline with custom-made software (A.S.) written in IgorPro 6.37 (Wavemetrics, Lake Oswego, OR). All recordings were performed at RT.

#### MEA recordings

We prepared acute hippocampal slices from 4-9 week old mice, and transferred them to a multi-electrode array (MEA) system (Cat# 60MEA200/30iR-Ti-gr; Harvard Apparatus, Holliston, MA), where they were perfused with carbogenated ACSF at a rate of at ∼2.5 ml/min. AAV transfection in Sham and Aβ42 mice was confirmed using an inverted fluorescence microscope (EVOS FL Color Cat# AMEFC4300; ThermoFisher Scientific, Waltham, MA) equipped with an EGFP filter set (Cat# AMEP4951; ThermoFisher Scientific, Waltham, MA). The MEA was composed of one stimulating, one ground, and 59 recording electrodes arranged in an 8×8 grid (electrode ⌀ = 30 µm; electrode spacing = 200 µm). We used constant current biphasic pulses (0.5 mA, 40 µs/phase, 80 µs total duration) to stimulate Schaffer collaterals and evoke subthreshold field potentials in CA1 *s.r.*. Extracellular voltage recordings were bandpass filtered at 1 Hz-3.3 kHz and sampled at 10 kHz via the Multi Channel Experimenter v2.20.15 software (Harvard Apparatus, Holliston, MA). After recording 20 sweeps in baseline conditions (i.e., no drugs), we added tetrodotoxin (1 µM) to the perfusing solution. Raw traces were exported from the Multi Channel Analyzer Software v2.20.15 as text files and analyzed with custom-made software written in MATLAB R2024a (A.K.M.). The analysis code allowed us to average 20 sweeps in baseline conditions and 20 sweeps in TTX. The average in TTX was subtracted from the average in baseline conditions to remove the stimulus artifact. We performed the FV and PSP analysis on recordings collected from the 2–3 recording sites in CA1 *s.r..* This analysis was performed using custom-made software (A.S.) written in IgorPro (Wavemetrics, Lake Oswego, OR). The TTX subtracted recordings were also used to generate videos. Here, we interpolated the recording from each MEA recording electrode using the MATLAB imgaussfilt function (σ = 0.2) and displayed it as a 50×50 pixel array. To calculate the spatial spread of the fEPSPs, we identified the maximal value of the fEPSP amplitude 10 ms after stimulating the Schaffer collaterals, and thresholded the pixel intensity as 1/e of this value. The area represented the number of pixels above this threshold value over the course of the entire recording. The charge transfer of the fEPSP was calculated between the time of the fEPSP onset and the time when the fEPSP amplitude was 0 µV. The charge transfer of the fIPSP was calculated between this point and the end of the recorded sweep, 50 ms after the stimulus artifact. The centroid of the fEPSP was calculated in a time window corresponding to 0.5 of the fEPSP peak amplitude, before and after its onset, as follows:

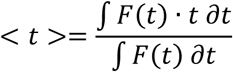

#### Histology and confocal imaging

Mice were deeply anesthetized with an intraperitoneal injection of pentobarbital (4 mg/g, w/w; catalog #54925-045-10; Med-Pharmex, Pomona, CA) and transcardially perfused with 20 ml of PBS 0.1 M and 20 ml of 4% paraformaldehyde in PBS (4% PFA/PBS) at 4°C. The dissected brains were postfixed overnight at 4°C in 4% PFA/PBS and cryoprotected at 4°C in 30% sucrose/PBS. Coronal sections (50 μm thick) were prepared using a vibrating blade microtome (VT1200S; Leica Microsystems, Deerfield, IL). All sections were postfixed for 20 min at 4°C in 4% PFA/PBS. The sections were then blocked and permeabilized for 1 h at room temperature (RT) in a solution of PBS containing BSA 3%, and Triton X-100 0.5%. The primary antibody incubation was performed by incubating the sections overnight at 4°C in a solution of PBS containing BSA 1%, and Triton X-100 0.3% and one or more of the following primary antibodies: guinea pig anti-S100β antibody (1:500; Cat# 287 004, Synaptic Systems, Göttingen, Germany; RRID:AB_2620025); guinea pig anti-GLT1/EAAT2 (1:500; Cat# AB1783, Millipore Sigma, St. Louis, MO; RRID:AB_90949); rabbit anti-GLT1/EAAT2 (1:1,000; Cat# NBP1-59632, Novus Biologicals, Centennial, CO; RRID:AB_11005463); rabbit anti-β-amyloid (1:500; Cat# 700254, Fisher Scientific, Waltham, MA; RRID:AB_2532306); rabbit anti-cleaved caspase-3 (Asp175) (5A1E) (1:1,000; Cat# 9664S, Cell Signaling Technology, Danvers, MA; RRID:AB_2070042); rabbit anti-K+/Cl-cotransporter (KCC2) (1:500; Cat# 07-432, Millipore Sigma, St. Louis, MO; RRID:AB_310611); rabbit anti-phospho tau (Ser 262) (1:500; Cat# PA5-85654, Fisher Scientific, Waltham, MA; RRID:AB_2792793). The secondary antibody incubation (see **Key Resource Table**) was performed for 3 h at RT using secondary antibodies diluted to 1:1,000 in 0.1% Triton X-100/PBS. The brain sections were mounted onto microscope slides using DAPI Fluoromount-G (catalog #0100–20; Southern Biotech, Birmingham, AL). For biocytin fills, biocytin 0.2%-0.4% (w/v) was added to the intracellular solution used to patch CA1-PCs and astrocytes. Each cell was filled for at least 20 min. The slices were then fixed overnight at 4°C in 4% PFA/PBS, cryo-protected in 30% sucrose PBS, and incubated in 0.1% streptavidin-Alexa Fluor 647 conjugate and 0.1% Triton X-100 for 3 hr at RT. The slices were then mounted onto microscope slides using Fluoromount-G or DAPI Fluoromount-G mounting medium (SouthernBiotech, Birmingham, AL). Confocal images were acquired using a Zeiss LSM710 and LSM980 inverted microscopes equipped with 488 nm Ar or 633 nm HeNe laser. All images were acquired as stitched z-stacks of 4 frames averages (1024×1024 pixels; 1 µm z-step) using a 40×/1.4 NA or 63×/1.4NA Plan-Apochromat oil-immersion objectives. Confocal images for spine analysis were also collected as z-stacks of 8 frame averages (1,024×1,024 pixels; 0.5 µm z-step; 3–5 digital zoom) using a 63×/1.4 NA Plan-Apochromat oil-immersion objective. Slices used for caspase staining under hypoxia conditions were maintained for 30 min without the carbogen mix (95% O_2_, 5% CO_2_) before being fixed with 4% PFA/PBS.

#### Dot blot experiments

Dot blot experiments were performed on protein extracts from the hippocampus of naïve WT mice, and WT mice that received AB-AAV hippocampal injections. The latter mouse cohort was injected with AB-AAV at P14-16, and tissue was collected after 3-6 weeks, when mice were roughly 2 month old. We used mice of either sex. Mice aged 18 and 24 months were obtained through the NIH/NIA Rodent Ordering System. Membrane and cytoplasmic protein extracts were obtained using the Mem-PER Plus Membrane Protein Extraction Kit (Cat# 89842; Thermo Fisher Scientific, Waltham, MA) according to the manufacturer’s instructions using a mixture of protease and phosphatase inhibitors (10 µl/ml, Cat# 78441; Thermo Fisher Scientific, Waltham, MA). The membrane protein extracts were used to measure protein levels of the glutamate transporters GLT-1 and GLAST. The protein concentration was determined using a Bradford assay (Cat# 5000006; Bio-Rad, Hercules, CA) and spectrophotometer measures. A standard curve for the Bradford Assay was produced using 2 mg/ml vials of BSA in PBS (Cat#23209; Thermo Fisher Scientific, Waltham, MA), with proteins are diluted to 0.25 mg/ml. Proteins from all mouse groups were directly spotted onto PVDF membranes and left to air dry (Cat# P2563; Millipore Sigma, Burlington, MA). The membranes were then blocked with 5% nonfat milk in TBST, pH 7.6, and probed using a primary antibody solution in which milk was replaced by BSA (5% BSA in TBST; pH 7.6). We used the following primary antibodies: rabbit anti GLAST (1:1,000, Cat# ab41751; Abcam, Cambridge, UK); rabbit anti GLT-1 (1:1,000, Cat# NBP1-59632; Novus Biologicals, Centennial, CO). The membranes were incubated with the primary antibodies overnight at 4°C. The secondary antibody incubation (biotinylated horse anti-rabbit IgG, Cat# BA-1100-1.5; Vector Laboratories) was performed for 1-2 h at RT with 5% nonfat milk in TBST, pH 7.6 at a dilution of 1:1,000. Control experiments for GLT-1 were performed using cell lysates from GLT-1^-/-^ mice, and no signal was detected (data not shown). We amplified the immunolabeling reactions with the Vectastain ABC kit (1:2,000; Cat# PK-6100; Vector Laboratories, Newark, CA) and the Clarity Western ECL (Cat# 170–5060; Bio-Rad, Hercules, CA) as a substrate for the peroxidase enzyme. For semiquantitative analysis, dot images were collected as 16-bit images using a digital chemiluminescence imaging system (c300, Azure Biosystems) at different exposures (1-3 s). Each image was converted to an 8-bit image for image analysis, which was performed using Fiji software. Only images collected at exposure times that did not lead to pixel saturation were included in the analysis. The intensity of each band was calculated as the mean gray value in an ROI surrounding each band of interest in three images collected using different exposure times.

#### Protein extraction from frozen hippocampus

For extraction of soluble Aβ_42_, chilled 100 µl RIPA buffer (1% NP-40, 0.5% sodium deoxycholate, 150 mM NaCl, 50 mM Tris hydrochloride, 0.5 mM MgSO_4_; Millipore Sigma; St. Louis, MO) with Complete Mini protease inhibitor (Millipore Sigma; St. Louis, MO) was added to frozen tissue. Samples were then sonicated on ice (6× pulses, 2×; separated by 3-5 min), and centrifuged for 30 min at 21,000 g at 4°C. Supernatants containing RIPA-soluble proteins, including soluble Aβ_42_ were pipetted off into new tubes. The remaining pellet was washed with 50 µL RIPA buffer with Complete Mini, and centrifuged a second time for 15 min at 21,000 g at 4°C. The RIPA-buffer containing supernatant was then combined with the first RIPA-buffer containing supernatant. The remaining pellet was then used for extraction of insoluble Aβ_42_. For that purpose, 150 µL chilled 5M guanidine–hydrochloride (Gu–HCl) buffer containing Complete Mini was added to the pellet, followed by vortexing for 2-3 s and sonication on ice (6× pulses). Centrifugation for 30 min at 21,000 g at 4 °C yielded a Gu-HCl soluble supernatant containing the insoluble Aβ_42_ fraction and other RIPA-insoluble proteins. RIPA and Gu-HCl supernatants were aliquoted and stored at −80°C. A BCA kit (Thermo Fisher; Waltham, MA) was used to determine total protein content for all fractions.

#### Quantification of Aβ_42_ in RIPA and Gu-HCl supernatants

Aβ_42_ was quantified using the Meso Scale Discovery K15200E V-PLEX kit (Meso Scale Diagnostics; Rockville MD) according to the manufacturer’s instructions. We used 2,000 ng of total protein from the RIPA-fraction per well for soluble Aβ_42_ and 1,000 ng of total protein from the Gu-HCl-fraction per well for insoluble Aβ_42_.

#### Open field test

In the open-field test, we monitored the position of a mouse freely moving in a white Plexiglas box (L=46 cm, W=46 cm, H=38 cm). Each mouse was video monitored for 15 min using a Live! Cam Sync HD webcam (Model #VF0770; Creative Labs). Videos were analyzed using EZTrack^109^.

#### Novel object location task

The OLT was used to evaluate spatial learning, which relies heavily on hippocampal activity^110^. C57BL/6J mice of either sex (6-10 week old) were handled for 5 min/day and acclimated to the empty behavioral arena (white Plexiglas box of W×L×H of 40×40×40 cm, uniformly illuminated at 45-49 lux) also for 5 min/day. Handling and habituation were repeated daily for 7 consecutive days. In the following training session (10 min), we positioned two identical objects in adjacent corners of the arena, 10 cm away from the edges. After 90 min, each mouse was subjected to a testing session (5 min), in which the two objects were positioned in opposite corners of the arena. The videos were analyzed using ezTrack v1.2, to calculate the amount of time spent within two 20×20 cm quadrants of the arena. The analysis was performed on 5 min of the habituation, training and testing sessions. The discrimination index (DI) was calculated as the time spent with the objected that was displaced (right quadrant, R), compared to the total amount of time spent in the proximity of the displaced and non-displaced object (left quadrant, L), according to the formula written below:

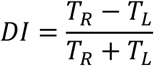

#### Data and statistical analysis

All experiments were conducted blind with regard to mouse genotype. Sample size determination was based on power analysis. Data averages are presented as mean±SEM, unless indicated otherwise.

Line profiles of KCC2 expressing cells was performed using FiJi. Briefly, we drew a line extending outward from the center of the soma of a CA1-PC, perpendicular to the plasma membrane. The length of this line was twice the distance from the soma center to the plasma membrane. For each cell, we generated one line profile and different line profiles were aligned at the location of the plasma membrane. Structural analysis of biocytin fills and dendritic spines were classified into four groups according to their neck and head size (i.e. mushroom, thin, stubby, filopodia), using Imaris 9.2^111–113^. Sholl analysis was performed using FiJi plugins. Functional analysis of astrocyte transporter currents were analyzed as described in previous works^48,52,53,86,114^. Briefly, we interleaved single and pairs of stimuli (100 ms apart), every 10 s. Single STCs were subtracted from paired STCs to isolate the current evoked by the second pulse. Single STCs were then shifted in time so that their onset matched that of paired STCs and subtracted from them. The resulting current represented the facilitated portion of the STC in response to paired stimuli (fSTC). Because the sustained potassium current superimposed to STCs does not facilitate, this method allows for almost perfect isolation of the STC, similarly to what can be achieved by subtracting the sustained potassium current in high concentrations of glutamate transporter antagonists from astrocytic currents recorded in control conditions. In the unlikely event that a proportion of the sustained potassium current was still present after subtraction, we approximated the rising phase of the residual current with the following mono-exponential function:

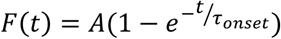

In our experiments, *τ_onset_* was set to 4 ms and A was scaled to match the amplitude of the residual potassium current, based on data collected from mouse astrocytes in our own previous works^48,114,115^. We used the fSTCs in control conditions and in the presence of a sub-saturating concentration of TFB-TBOA (1 µM) to calculate the time course of glutamate clearance by performing a deconvolution analysis of the STCs. *First*, the STCs recorded under control conditions and in TFB-TBOA (1 µM) were binomially smoothed and fitted with the following equation^116^:

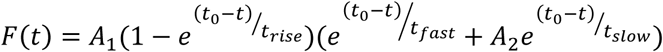

*Second*, we approximated glutamate clearance in TFB-TBOA (1 µM) with an instantaneously rising function decaying mono-exponentially, with the same time course of the fSTC. *Third*, we deconvolved this approximated glutamate clearance from the fSTC recorded in low TFB-TBOA to obtain the filter. *Fourth*, we deconvolved the filter from the fSTC recorded in control conditions to obtain the glutamate clearance waveform in control conditions^48,52,53,114,115^. Miniature events (mEPSCs and mIPSCs) were detected using an optimally scaled template adapted for Igor Pro (A. S.). PPRs were calculated by subtracting single from paired E/IPSCs averaged across 20 traces, after first peak normalization. Statistical analysis was performed using IgorPro 6.37 or IBM SPSS Statistics 28. Statistical significance was determined by Student’s paired or unpaired t-test or Mann-Whitney test, as appropriate or, when comparing multiple groups, by one- or two-way ANOVA, with or without repeated measures. A full report of all statistical comparisons for this manuscript is included in the data sheets shared via Open Science Framework. Differences were considered significant at p<0.05 (*p<0.05; **p<0.01; ***p<0.001).

