## Supplementary figures and images for "An increased excitation and inhibition onto CA1 pyramidal cells sets the path to Alzheimer’s disease"

### Supp. Fig. 1

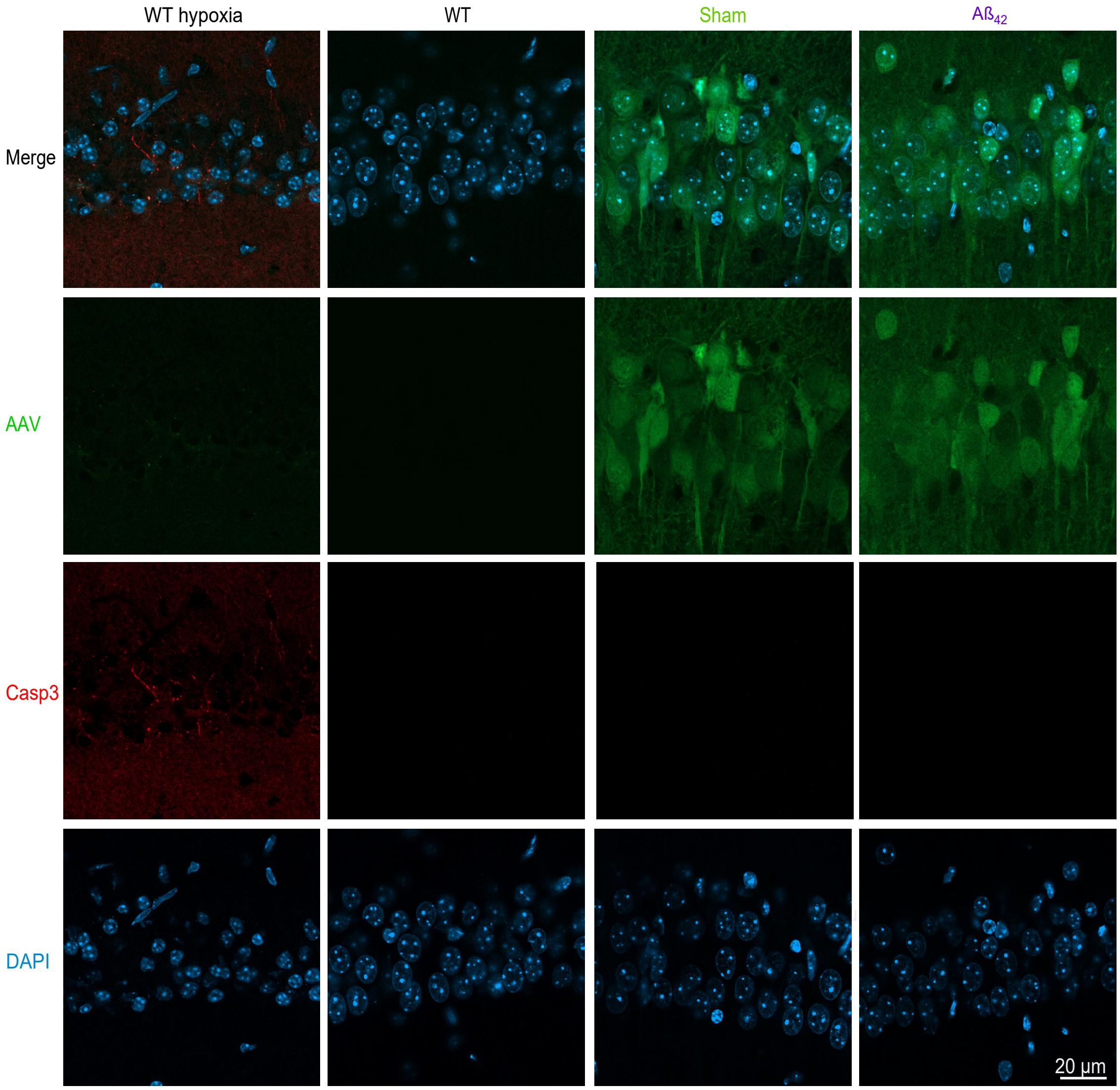


**Supplementary Figure 1**

### Supp. Fig. 2

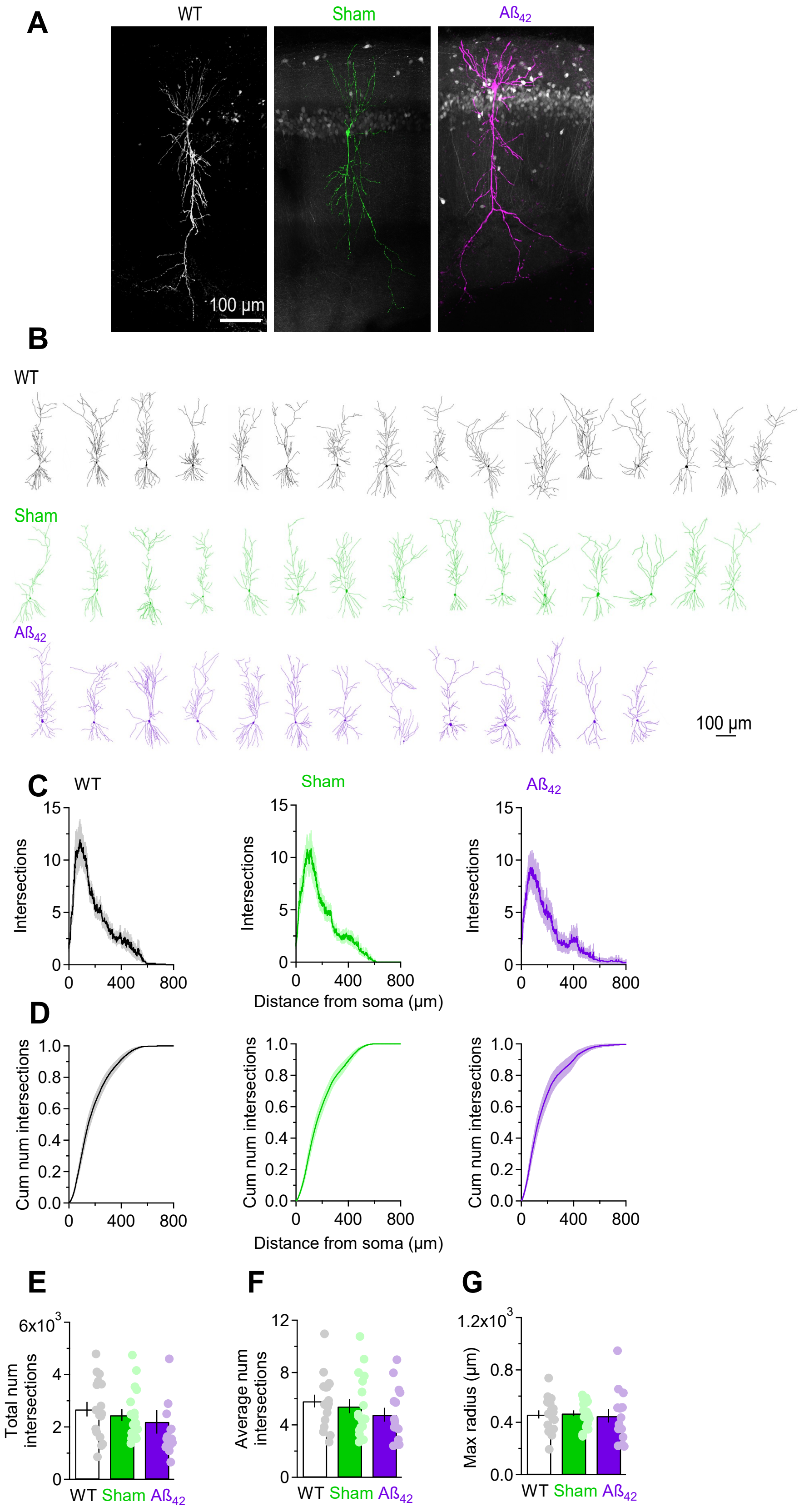


**Supplementary Figure 2**

### Supp. Fig. 3

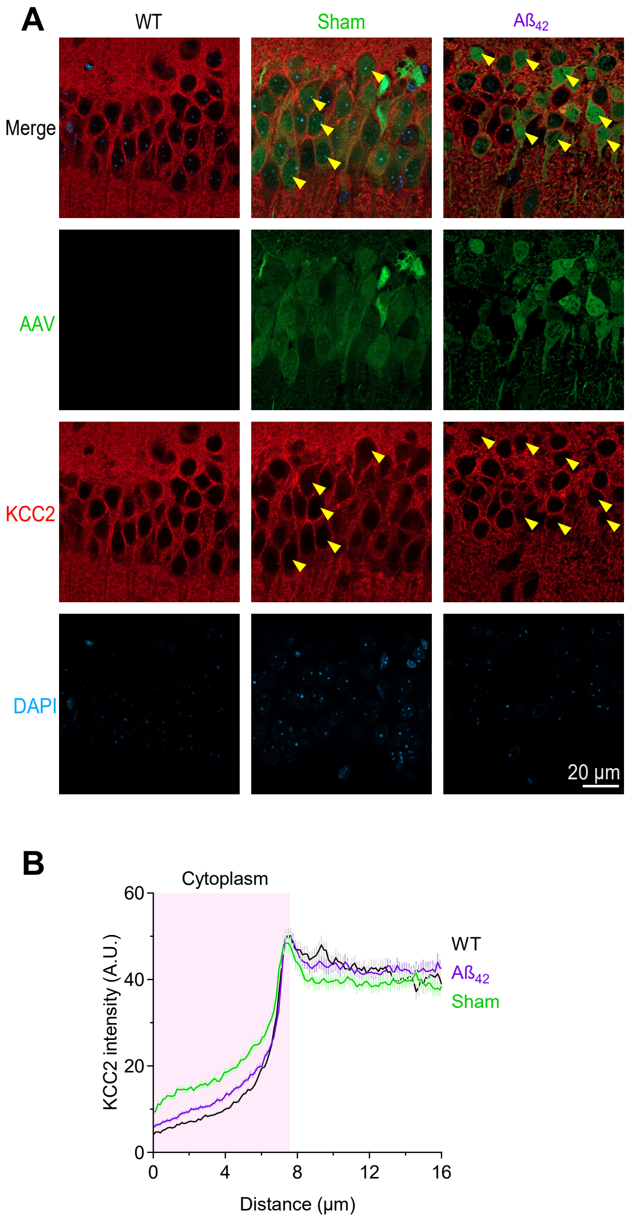


**Supplementary Figure 3**
